# Arterial arcades and collaterals regress under hemodynamics-based diameter adaptation: a computational and mathematical analysis

**DOI:** 10.1101/2024.07.22.604568

**Authors:** Vivi Rottschäfer, Willem G.N. Kuppers, Jiao Chen, Ed van Bavel

## Abstract

Segments in the arterial network have a >1000-fold span of radii. This is believed to result from adaptation of each segment to the wall shear stress (WSS), with outward respectively inward remodeling if WSS is higher or lower than some reference value. While this seems a straightforward mechanism for arterial tree design, the arterial network is not a tree but contains numerous arcades, collaterals and other looping structures. In this theoretical study, we analyzed stability of looping structures in arterial networks under WSS control.

Simulation models were based on very simple network topologies as well as on published human coronary and mouse cerebral arterial networks. Adaptation was implemented as a rate of change of structural radius of each segment that is proportional to the deviation from its reference WSS. A more generalized model was based on adaptation to a large range of other local hemodynamic stimuli, including velocity, flow and power dissipation.

For over 12,000 tested parameter sets, the simulations invariably predicted loss of loops due to regression of one or more of the segments. In the small networks, this was the case for both the WSS and the generalized model, and for a large range of initial conditions and model parameters. Loss of loopiness also was predicted by models that included direction-dependent adaptation rates, heterogeneous reference WSS or adaptation rates among the adapting segments, and adaptation under dynamic conditions. Loss of loops was also found in the coronary and cerebral artery networks subjected to adaptation to WSS.

In a mathematical analysis we proved that loss of loops is a direct consequence of Kirchhoff’s circuit law, which for each loop leads to a positive eigenvalue in the Jacobian matrix of partial derivatives in the adaptation model, and therefore to unstable equilibria in the presence of loops.

Loss of loops is an inherent property of arterial networks that adapt to local hemodynamics. Additional mechanisms are therefore needed to explain their presence, including direct communication between connected segments.

## Introduction

The arterial system includes a myriad of blood vessels, each with its own diameter, varying from ∼ 2 cm to ∼10 micrometer. Individual arterial segments obtain diameters that seem well-matched to the carried blood flow. This observation has inspired research on the design principles and their underlying mechanisms for centuries. A well-known design principle is ‘Murray’s law’ [1], which states that in individual segments within a network, the cube of the diameter is proportional to the flow. Murray based this principle on cost minimization, where costs include those of power dissipation and vascular volume. Much later, it became clear that adaptation of vascular structure and diameter, so-called arterial remodeling, in response to blood flow depends on endothelial cells [2]. Substantial evidence indicates that such remodeling involves the sensing of wall shear stress (WSS) by the endothelial cells. This evidence includes regulation of WSS by remodeling following experimentally induced changes in flow in the canine carotid artery [3], rat mesenteric arteries [4] and vessels in the chick chorioallantoic membrane [5]. WSS regulation by remodeling was also shown in isolated perfused porcine coronary arteries in organoid culture [6]. These structural responses of vessels to changes in flow typically take weeks.

In Poiseuille flow WSS is proportional to flow divided by diameter cubed. It thus seems that WSS sensing by endothelial cells underlies Murray’s law. A WSS higher than some reference level would induce an increase of arterial diameter by outward remodeling, while a WSS below the reference level would cause inward remodeling. This adaptation shapes arterial trees towards increasingly smaller diameters at each branching level, and reshapes the network in response to changes in flow. An increase in flow and outward remodeling of the network may result from organ growth during development and (for skeletal muscle) following physical exercise. A decrease in flow and WSS-driven inward remodeling could result from the development of a flow-limiting atherosclerotic plaque in an upstream main artery [7].

The arterial network is not merely a tree starting from the aorta and branching towards the billions of capillaries. Rather, many organs contain arterial arcades and so-called collateral arteries that provide alternative routes for blood flow if an artery becomes blocked. Fig. 1 provides examples.

**Fig. 1:**
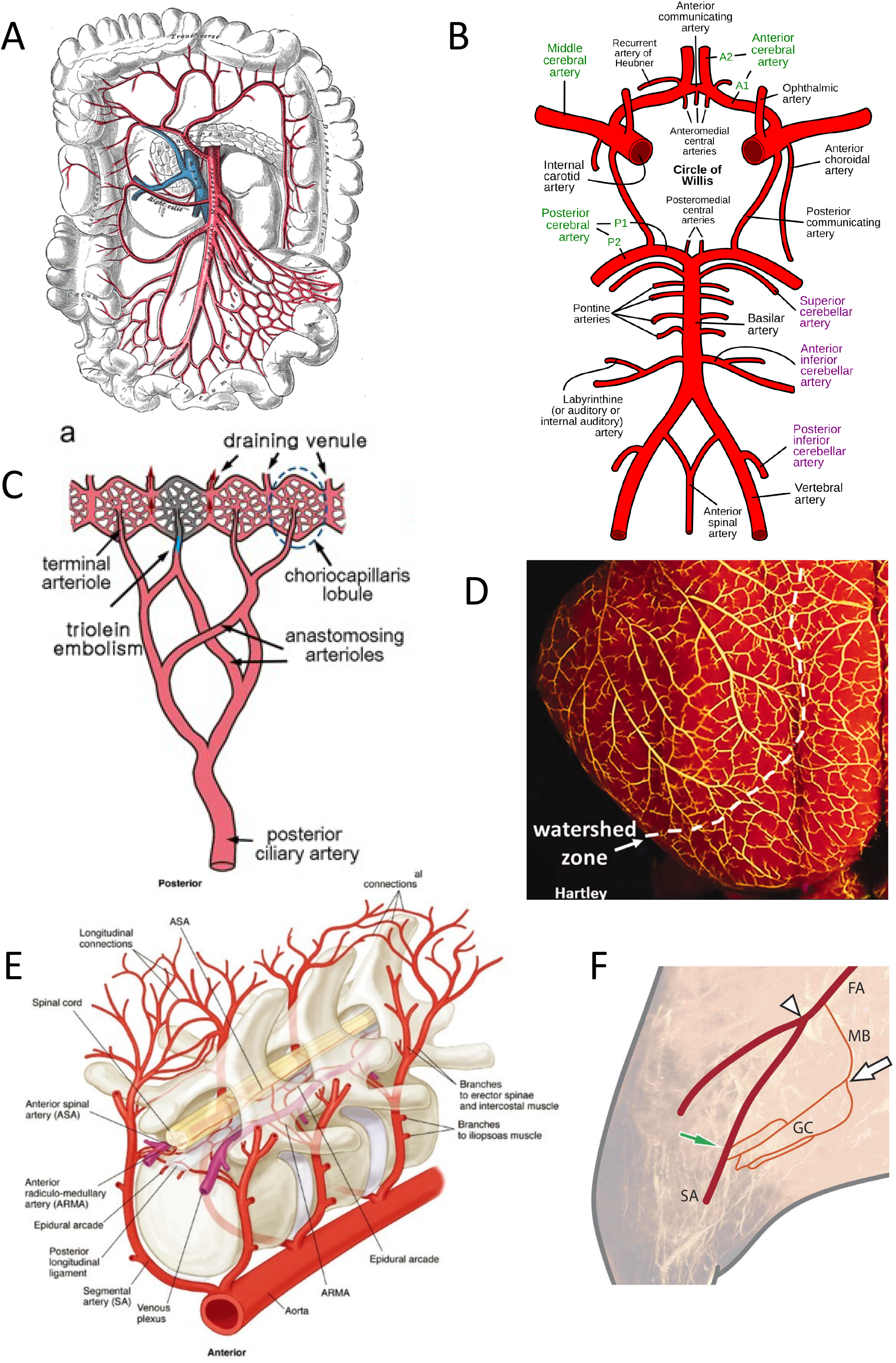
Examples of arterial collaterals and arcades. A: arcades in the mesenteric arterial bed that perfuses the intestine, reproduced from [40]. B: the Circle of Willis, reproduced from [41]. C: anastomosing arterioles in the choroid circulation that is located underneath the retina in the eye, reproduced from [42]. D: leptomeningeal collaterals on the surface of the guinea pig brain cortex, demonstrating collaterals both within and between perfusion territories, the latter crossing watershed zones. Reproduced from [43]. E: longitudinal and lateral arterial anastomoses in the spinal cord, reproduced from [44]. F: collaterals in the gracilis muscle (GC) in the upper leg, between the muscular branch (MB) of the femoral (FA) and the saphenous artery (SA), based on arterial reconstruction in the rat, reproduced from [45].

Arcades are loosely defined as ‘roundabouts’ of arteries of roughly similar size which are connected to multiple larger entrance vessels and smaller downstream vessels. The mesenteric arcading arteries (Fig. 1A) and the Circle of Willis in the brain (Fig. 1B) are clear examples of this. Much smaller anastomosing arterioles are found in the choroid circulation in the eye (Fig. 1C). The leptomeningeal arteries in the brain show extensive arcading connections within perfusion territories of the major cerebral arteries, as well as collateral connections between these territories (Fig. 1D). The arterial network in the spinal cord (Fig. 1E) forms an extensively looping system at multiple branching levels. Tiny collateral cross-connections exist between major arteries, e.g. in the leg between femoral and saphenous arteries (Fig. 1F). We will refer to all these topologies as ‘loops’.

The presence of loops in e.g. the heart and brain is variable between subjects, resulting in large differences in outcome when a major artery is blocked, as in acute myocardial infarction[8] and ischemic stroke [9]. The causes of such variability are largely unknown, and may relate to differences in adaptation to local hemodynamics and other stimuli.

In a network, reduction of the diameter of one of the segments may lead to either an increase or decrease of WSS in that segment. This depends on the distribution of resistances, rendering WSS-dependent adaptation a complex mixture of negative and positive feedback loops. Therefore, despite the current evidence for WSS-driven adaptation in arterial networks, this concept raises a concern on network stability. Already in 1996, Hacking et al. [10] mathematically derived that WSS-driven adaptation in a looping network of two parallel arteries leads to regression of at least one of the two vessels and thereby loss of the loop. In the same study, numerical simulation of adaptation to WSS in a honeycomb-like looping topology showed regression towards a single connection between the entrance and exit point.

While this concern has been appreciated for several decades, an extensive analysis of stability of loops in networks under WSS regulation is lacking. We hypothesize that arterial networks subjected to WSS-driven diameter regulation will always lose their loops, reducing these networks to mere arterial trees without any cross-connections. In the current work, we address this by extensive computational analysis of adaptation in small networks. We included several variations of adaptation models that are based on local hemodynamics. We further studied adaptation in very extensive arterial networks based on published coronary and cerebral arterial topologies. Finally, we provide a mathematical analysis of the instability of loops, also for several variations of adaptation models.

## Results

### Arterial loops regress in WSS-dependent vascular adaptation

Loop stability was first tested in simulations of adaptation to WSS in very small arterial networks. Simulations were based on the rule that radii of vessels adapt according to the WSS, combined with algebraic hemodynamic relations: the Poiseuille relation, Kirchhoff’s flow law, and the dependence of WSS on pressure gradient and radius. A dimensionless model with reduced parameter space was applied. Fig. 2 shows a triangular network of adapting arterial segments (colored), each with its conductance 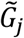 depending on time-varying radii 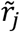 based on the Poiseuille relation. The network is connected to three external pressures 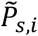, via source conductances 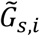.Adaptation of each segment was based on its absolute wall shear stress, 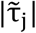, according to

**Fig. 2:**
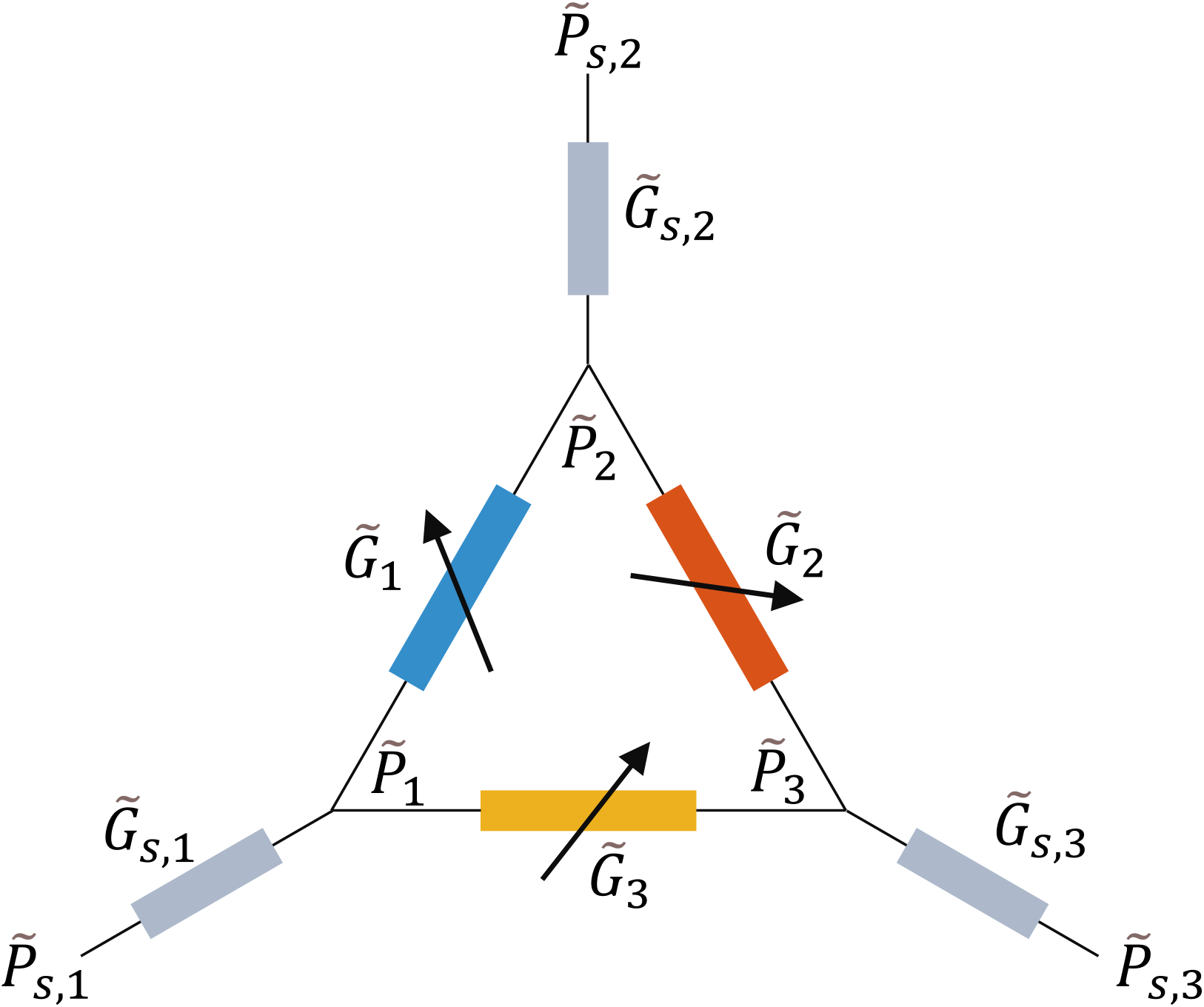
The triangular example network with definition of pressures and conductances, shown as electrical equivalent network. The three colored segments form a loop. Each of these segments is able to adjust its radius and therefore its conductance. The sources have a fixed source pressure and source conductance.

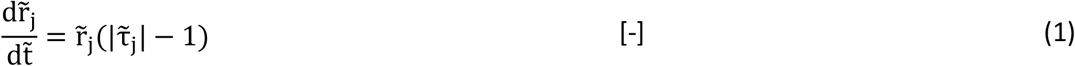

The tildes indicate dimensionless quantities, with the reference level for 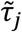 taken as 1. The methods section below provides further details. The model parameters are listed in Table S1. In this model the vessel radius is at equilibrium when the absolute WSS of all adapting segments equals this reference level 1, while a higher or lower WSS causes outward and inward remodeling, respectively. In turn the WSS in each segment depends on its radius and the pressures, causing a hemodynamic coupling of adaptation between the three segments.

Fig. 3 shows examples of adaptation in this single-loop triangle topology (Fig. 3A) as well as in a pentagon topology with six adapting segments and a double loop (Fig. 3B). In these examples, sets of non-zero radii could be found where WSS in all segments were at reference. In the simulations, the initial radii were set 1% above these equilibrium radii. All radii initially returned to these equilibria, but then, around t=10, one (triangle) and two (pentagon) segments, respectively, fully regressed, resulting in a loss of the single loop (triangle) and double loop (pentagon) and a transition to a tree structure. The transition was associated with large changes of WSS, while in steady state WSS of remaining segments had returned to reference and the WSS of regressed segments fell to zero. These transitions to trees resulted in a new stable configuration for the remaining segments. These example simulations suggest that, if an equilibrium exists where all segments in a looping network are at reference WSS, this equilibrium is unstable under WSS-driven adaptation of vascular radius, and a minimal deviation from this equilibrium causes the system to progress to a non-looping network with regulated WSS.

**Fig. 3:**
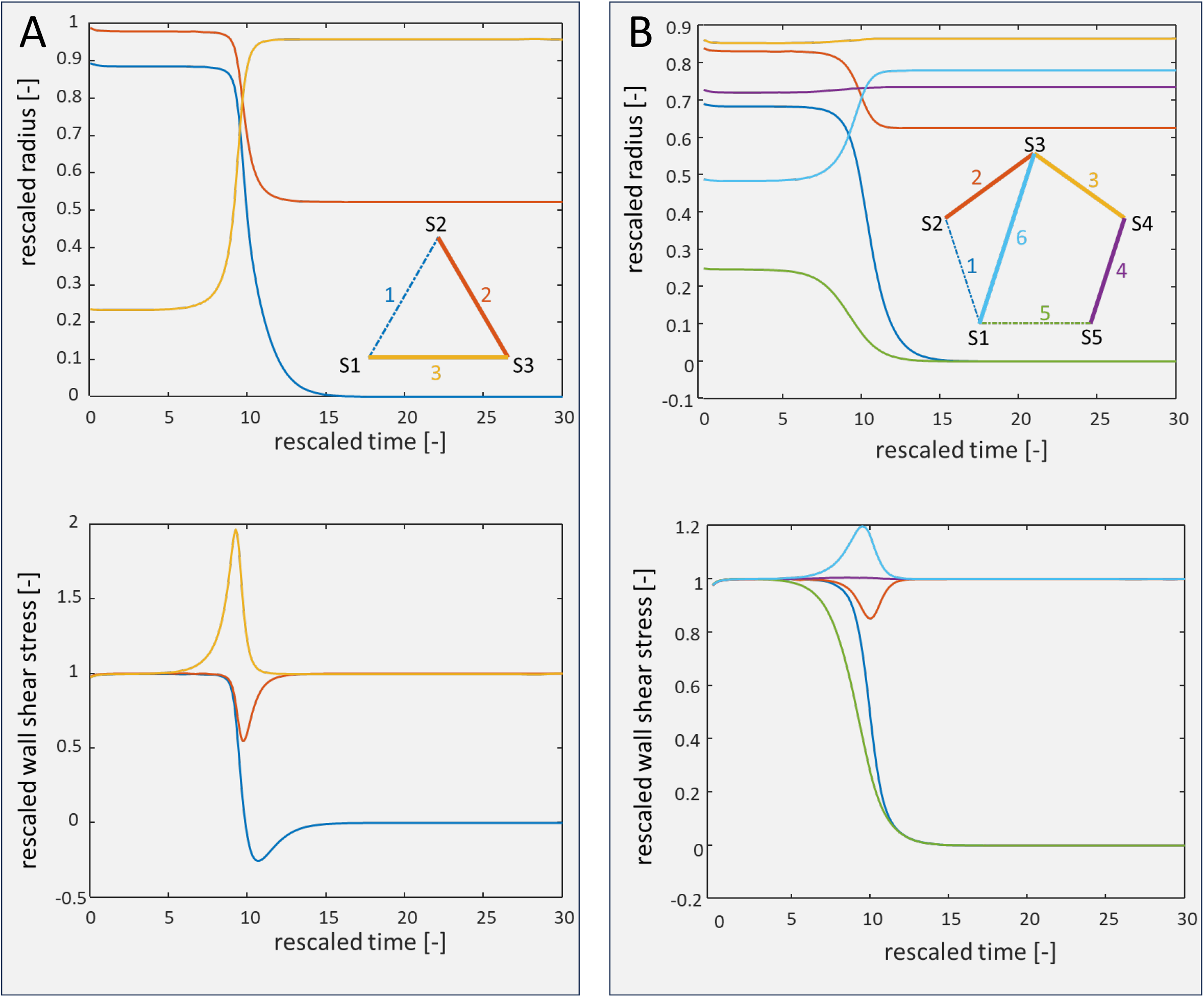
WSS-dependent adaption in example networks (see Table S1 for parameter values). Simulation started at 1.01 times the radii for equilibrium WSS based on a simplex search. Top: rescaled radius. Bottom: rescaled WSS. See methods for rescaling to dimensionless quantities. A: triangle topology, demonstrating regression of segment #1 (blue) and thereby loss of the loop. B: pentagon topology with a cross-connection, showing regression of segment #1 (blue) and #5 (green), and thereby loss of both loops.

While the above simulations started close to an unstable equilibrium, loss of loopiness was also found for a very large set of non-equilibrium initial radii. The initial conditions determined which and how many segments regressed (Tables 1 and 2). For the triangle example, there are 2^3^=8 possible combinations of stable and regressed segments. Six such combinations were found for this example, with 0-2 but not 3 segments remaining. For the pentagon, 7 of 64 possible combinations were found, with 3 or 4 segments remaining. No combinations were found that left a loop intact (i.e. with remaining segments 1-2-6, 3-4-5-6 or 1-2-3-4-5). For very small initial radii, such that all WSS started below the reference values, the full network regressed (not shown).

**Table 1:** Triangle model (see Fig. 3A). Steady state rescaled radii as observed during simulations with 9^3^= 729 combinations of initial radii, in the range 0.05x to 4x the initial unstable equilibrium values. Shown are six observed sets of final radii, the number of remaining segments and loops in these sets, and the number of simulations (i.e. combinations of initial radii) in which each set of final radii was observed. None of the simulations resulted in three remaining segments and a remaining loop.

| Steady state normalized radii [-] |  |  | Number of remaining segments | Number of remaining loops | Found number of simulations |
| --- | --- | --- | --- | --- | --- |
| $\tilde{r}_1$ | $\tilde{r}_2$ | $\tilde{r}_3$ | | | |
| $> 0$ | $> 0$ | $> 0$ | 3 | 1 | 0 |
| 0 | 0.522 | 0.959 | 2 | 0 | 202 |
| 0.445 | 0 | 0.998 | 2 | 0 | 109 |
| 0.892 | 0.984 | 0 | 2 | 0 | 252 |
| 0 | 0 | 0.981 | 1 | 0 | 76 |
| 0 | 0.764 | 0 | 1 | 0 | 48 |
| 0 | 0 | 0 | 0 | 0 | 42 |

**Table 2:** Pentagon model (see Fig. 3B). Sets of remaining segments as observed during simulations with 3^6^ = 729 combinations of initial radii, in the range 0.3x to 3x the initial unstable equilibrium values. Of 64 possible combinations of remaining segments, 7 combinations were observed. None of the simulations resulted in remaining loops.

| Remaining segments<br>(for numbering see Fig. 3B insert) | Number of remaining segments | Number of remaining loops | Found number of simulations |
| --- | --- | --- | --- |
| 1 2 3 4 5 6 | 6 | 2 | 0 |
| Any five of the six segments | 5 | 1 | 0 |
| 2 3 4 6 | 4 | 0 | 198 |
| 1 3 5 6 | 4 | 0 | 11 |
| 1 3 4 6 | 4 | 0 | 15 |
| 1 2 3 4 | 4 | 0 | 144 |
| 1 3 6 | 3 | 0 | 11 |
| 1 4 6 | 3 | 0 | 349 |
| 2 3 6 | 3 | 0 | 1 |

The loss of loopiness as a result of WSS-dependent adaptation was invariably found for a wide variety of model parameters and initial radii. Fig. 4 and Table S2 show these data for the triangle example (A) and pentagon model (B). For each set of parameters, we first attempted to find equilibrium radii where WSS of all three segments equaled their reference value. The top panels of Fig. 4 illustrate that such radii indeed exist for the vast majority of parameter values. We then simulated the models, starting from initial radii that were 0.3x to 3x those of the top panel. The bottom panel indicates the number of segments that remained after network adaptation and that obtained reference WSS. For the triangle (Fig. 4A), two segments remained for the majority of the combinations of the parameters (n=563 of 729 models). Less frequently, one segment remained (n=134) or all segments were lost (n=42). In none of the cases, all three segments remained. For the pentagon (Fig. 4B), in most simulations 3 (n=984 of 3072 models) or 4 segments (n=2051) remained, but simulations never resulted in 5 or 6 remaining segments. For both topologies and all parameters, loops disappeared.

**Fig. 4:**
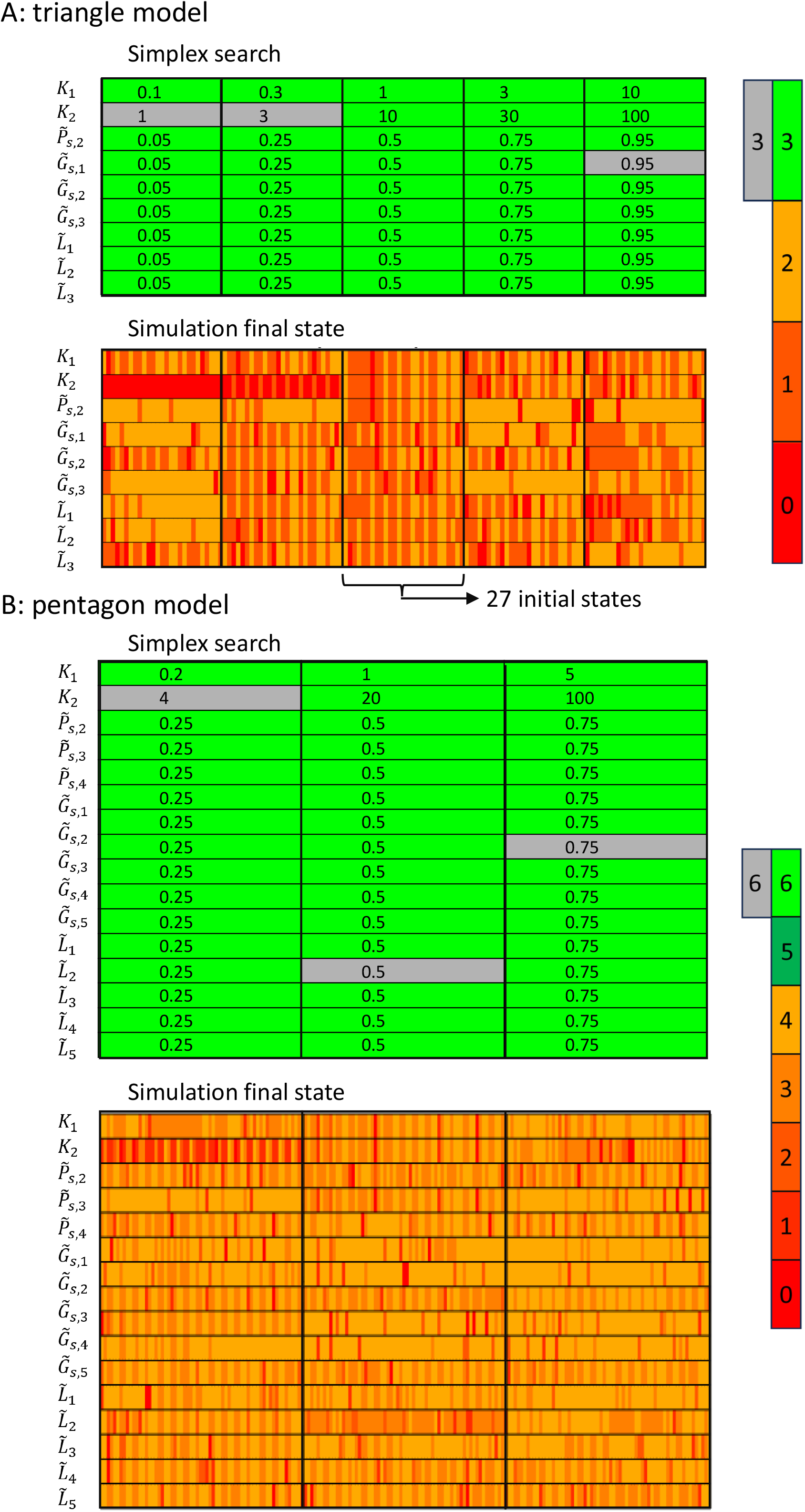
Effect of variation of parameters and initial states for the triangle model (A) and the pentagon model (B). Top panels in A and B: result of the simplex searches. Indicated are the varied parameter and their values. For variations of rescaled source conductance 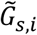 for the indicated sources, the values for the other sources were also adapted in order to maintain 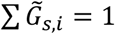. This was also done for the indicated variations of segment length 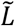. Green: Cases where the simplex search provided an (unstable) equilibrium. Gray: cases where this was not the case. Bottom panels: fingerprint results of the simulation of adaptation to WSS. Each vertical line is a simulation. For each parameter set, a range of initial states was applied. For the triangle model, these were all combinations of 0.3, 1.01 and 3 times the results of the simplex search (27 simulations per parameter set, in total 1215 simulations). For the pentagon, initial states were combinations of 1.01 and 3 times the simplex search results (64 simulations per parameter set, in total 3072 simulations). In all simulations, all segments had either regressed or had achieved WSS*ref* in the final state. Colors indicate the number of stable segments, as defined by the color bars on the right. At most two (triangle) or four segments (pentagon) remained, and none of the simulations resulted in remaining loops.

### Loss of loopiness by local hemodynamic adaptation occurs in a wide variety of models

Since stable arterial arcades exist in real life, we sought for alternative adaptation models that do predict presence of loops in adapted networks. Table S2 provides details on the models, parameters and predictions. In all these alternatives, adaptation occurs to a local hemodynamic stimulus.

#### Generalized adaptation to hemodynamic conditions

We considered a range of alternative hemodynamic stimuli for adaptation, by applying a more generalized adaptation model:

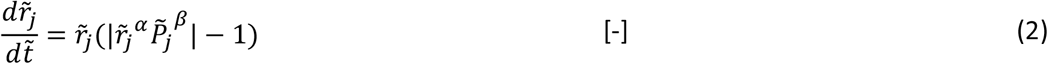

With parameters *α*>*0 and β*>*0*. For *α=1* and *β=1* this reduces to the above model of WSS regulation. Choosing *α=2* and *β=1* reflects regulation of blood flow velocity. *α=4* and *β=1* would indicate regulation of flow. Taking *α=4* and *β=2* reflects regulation of power dissipation. The simulation results reveal loss of loopiness in all tested cases for a range of *α* and *β* values, including these specific values (Table S2).

#### Direction-dependent adaptation to wall shear stress

The biological processes involved in inward and outward remodeling may be different, leading to different rates of adaptation during shrinkage and growth. This was modeled by a direction-dependent rate of adaptation to WSS. We tested the effect of a 5x faster or slower outgrowth rate. While in some cases this affected the final topology, no loops remained (Table S2).

#### Different reference WSS (WSSref) or adaptation speed among adapting segments

Arteries are sensitive to a myriad of chemical and physical stimuli, which may differ between segments. Additive effects of such stimuli and WSS on adaptation can be modeled by different reference WSS (WSS*ref*) among the adapting segments. In addition, such stimuli may affect adaptation speed. As shown in Table S2, again no remaining loops were observed in models with heterogeneous reference WSS or adaptation speed.

#### Adaptation under dynamic conditions

The circulation is a dynamic system: systemic pressure is pulsatile due to the heart beat, and to some extent, the respiratory cycle, but also due to periods of activity, postural changes, and the diurnal rhythm. Such dynamics cause fluctuating flow in all segments in a loop, including possible flow reversal. It seemed conceivable that this leads to sufficient stimulation of the adaptation process to obtain loop stability, especially since we took the absolute value of WSS as the stimulus. Fig. 5 provides examples of adaptation during sine wave oscillations of parameters. In Fig. 5A, source conductance 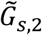 (see Fig. 2) was varied in the example triangle topology of Fig. 3. In Fig. 5B, source pressure 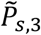 was varied in the example pentagon model. Such oscillations reflect rhythmic changes in vasoconstriction in the networks forming the sources and sinks of the adapting models. We tested oscillation frequencies that were much slower (left panels), in the same order (middle), or much faster (right) than the adaptation dynamics, with amplitudes equal to 50% of the mean value. As can be seen, loop regression also occurred under this dynamic regime. For both examples, the same segments regressed for all three frequencies. The remaining segments showed substantial oscillations in radius at low but not high frequency. Reversely, oscillations in WSS around the reference value were small at low frequency and large at high frequency. This reflects the ability of this network to regulate WSS during slow oscillations better than during fast oscillations.

**Fig. 5:**
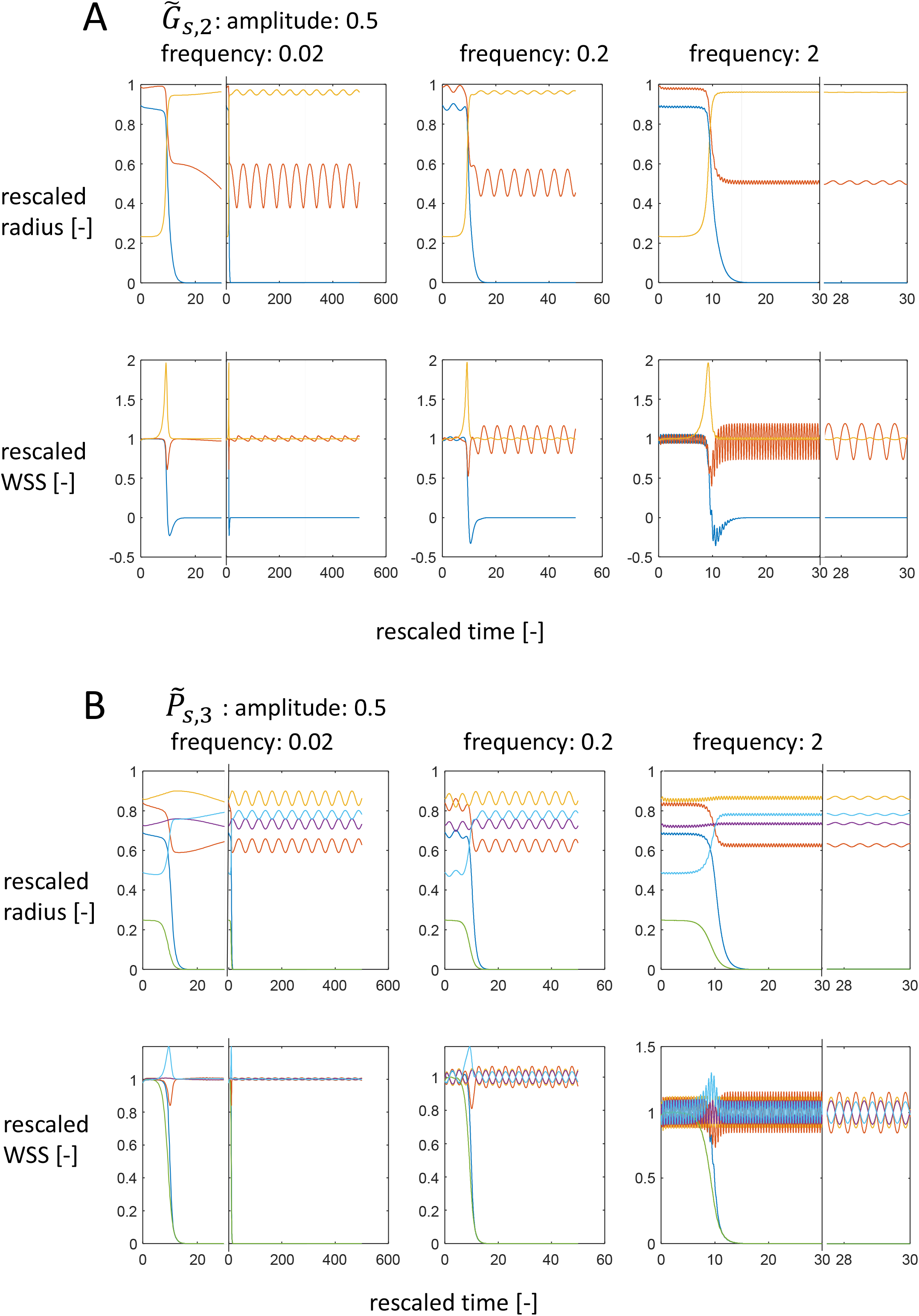
Examples of WSS regulation under dynamic conditions. A: triangle model with sine-wave oscillations of the second rescaled source conductance 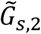 around the default value of 0.33. B: pentagon model with oscillating third rescaled source pressure 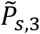 around the default value of 0.4. Panels indicate rescaled radius and WSS for rescaled frequencies of 0.02 (including a zoom-in of the adaptation transient), 0.2 and 2 (including a zoom-in of oscillations after the adaptation transient).

We considered oscillations of all parameters with the exception of segment lengths. This included dynamics in systemic pressure and in balances of source and network conductances. Loss of loopiness was observed for all simulations under dynamic conditions, as summarized in Table S2. Final topology was not always equal for the three frequencies because the high amplitude may bring the state to the domain of a different attractor, notably at low frequencies.

### Adaptation in experimentally observed large arterial networks

Also in large arterial network models based on experimental data from the human coronary and mouse cerebral circulations, adaptation to WSS caused regression of loops. For the human coronary circulation, data from Schwarz et al. [11] were used. In brief, these authors filled the arterial bed of a human heart with fluorescent plastic and cut the heart by a so-called imaging cryomicrotome [12].

Fluorescence images were taken of the remaining block after each cut, and these images were combined to 3D stacks. These authors then reconstructed the 3D images, segmented the arterial bed and translated this into a network description of arterial segments and nodes. Segments down to 15 µm in radius were included. In order to calculate network hemodynamics, the authors estimated conductance of the downstream pathways by extrapolation of the observed branching characteristics.

This coronary network consisted of 101,223 segments with two entrances and 100,961 connections to the downstream pathways. The network contained 3202 loops. We implemented adaptation of arterial radius to WSS in this coronary network. Fig. 6A shows the adaptation dynamics for WSS*ref*=20 Pa. As can be seen, all loops rapidly disappeared (blue line in top panel), between day 4 and day 21. In addition, at a slower pace, almost half of the segments and outflow connections regressed. Meanwhile, coronary inflow increased from 1.14 L/min to 2.52 L/min, indicating higher conductance due to increased diameters of the remaining segments. These flows are unrealistically high. We therefore decreased the downstream conductances to respectively 50% (6B) and 25 % (6C) of the published values. This provided more realistic flows, but further loss of segments.

**Fig. 6:**
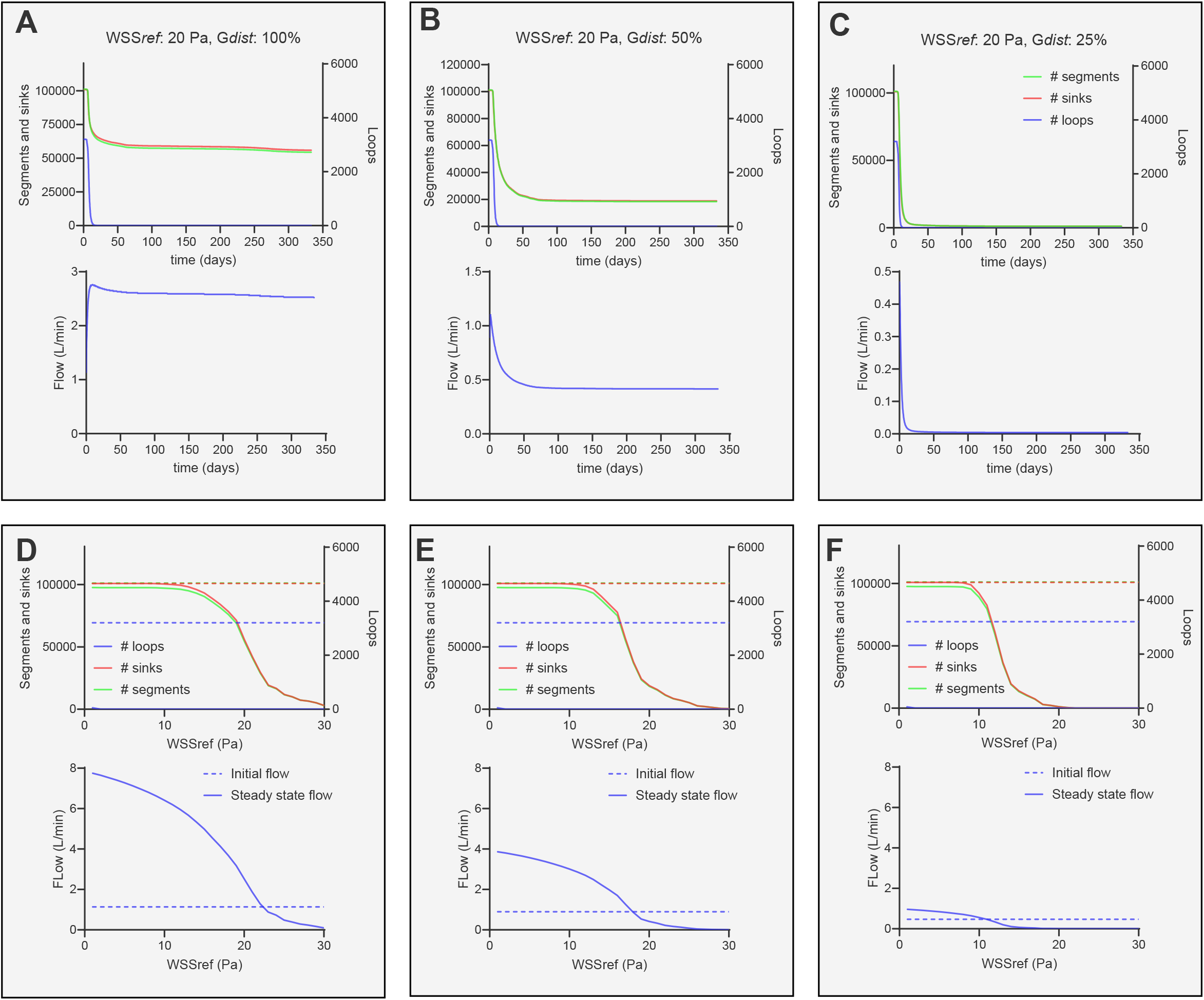
Adaptation to WSS in the human coronary network. 6A-C: adaptation dynamics based on WSS*ref*=20 Pa and taking G_dist_ equal to 100%, 50% and 25% of the published values respectively. Indicated in the upper panels are the number of segments (green), sink connections (red) and loops (blue, right vertical axis scale). The lower panels show the associated change in coronary inflow. 6D-F: Steady state results for WSS*ref* between 1 and 30 Pa and G_dist_ equal to 100%, 50% and 25% of the published values respectively. Upper panels show initial (dashed lines) and steady state (solid lines) number of segments, sinks and loops. Lower panels show initial (dashed lines) and steady state flow (solid lines) as a function of WSS*ref*.

We tested the effect of WSS*ref* on this adaptation. The supporting movie visualizes the steady state results for increasing WSS*ref* in each frame. At low WSS*ref* (first frames), regression is seen in predominantly sub-endocardial layers, indicating loss of the loops that were present here. Above WSS*ref*= 10 Pa, more and more segments regressed, notably in the sub-endocardium. The regression occurred clustered in specific areas of the left ventricle.

Figs 6D-F show the steady state results of these simulations for WSS*ref* between 1-30 Pa and distal conductances of 100%, 50% and 25% of the published values, respectively. Loops again disappeared in all simulations. Only for WSS*ref* = 1 Pa, a few loops were still present at the end of the simulation. This is a result of the fact that the simulation had not reached full steady state yet. For low WSS*ref*, ∼ 3400 segments regressed (solid versus dashed green lines). This corresponds to the regression of the 3202 loops, while almost all connections to the downstream circulation (red lines) remained intact. Above ∼ 9-13 Pa, more and more segments regressed, leading to an increasing loss of outflow connections for higher WSS*ref*. This loss of outflow connections was most prominent for simulations based on 25% of the published downstream conductances (6F).

We also simulated network adaptation based on published cerebral artery network data of nine mice [13, 14]. These data were based on optically cleared brain tissue imaged by lightsheet microscopy, followed by extensive image analysis, and contained in the order of 10^6^ – 10^7^ segments. We built several smaller arterial networks from these data by replacing the very many smallest arterioles and capillaries by lumped non-adapting downstream conductances. We kept ‘coarse’ and ‘fine’ networks of respectively ∼10^4^ and ∼10^5^ segments (see methods). Fig. 7A-B demonstrates the dynamics of adaptation for a BALBc mouse with a ‘fine’ segment selection at WSS*ref*=5 Pa and 20 Pa, respectively. In these models, loop regression started after approximately 100 days, and eventually all loops again regressed. At 5 Pa (7A), while segments that make up loops regressed, no sink connections disappeared, and flow increased. At 20 Pa (7B), also the vast majority of segments and sink connections disappeared, and flow became very small. As was the case for the human coronary bed, adaptation caused a loss of all loops for all WSS*ref*. Outflow pathways connecting to the capillary bed disappeared more and more for higher WSS*ref* (7C). We determined the WSS*ref* levels that caused at least 50% reduction of loops, segments, sinks and flow in nine mice (3 BALBc mice, 3 C57BL/6 mice, 3 CD1-E mice). Loops disappeared for any WSS*ref* and for both ‘coarse’ and ‘fine’ networks in all mice. Regression of segments and sink connections occurred above 12 to 17 Pa, with slightly lower values for the BALBc mice compared to the other two strains (7D-E).

**Fig. 7:**
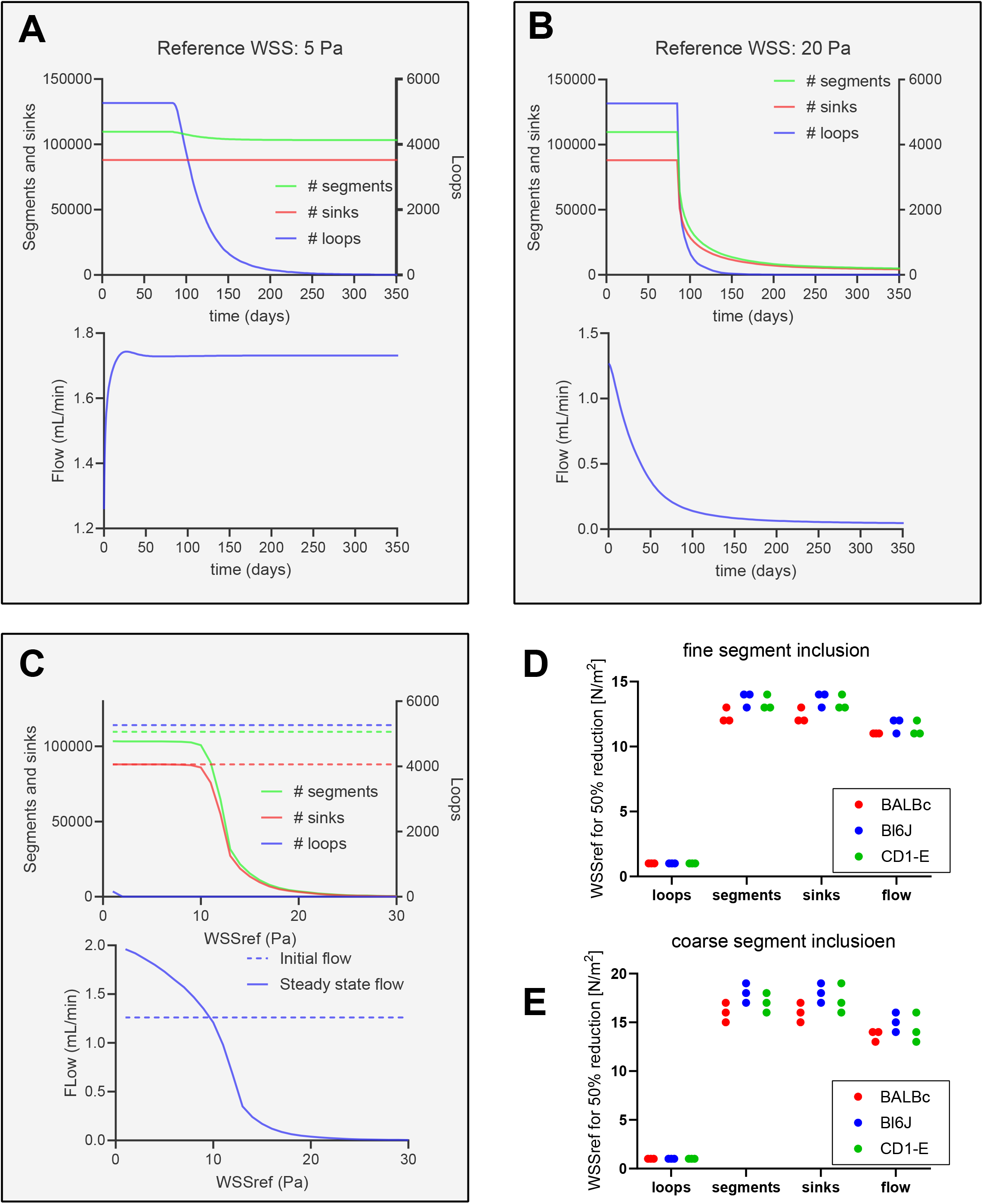
Adaptation to WSS in the cerebral artery networks. 7A: adaptation dynamics, based on a BALBc mouse with ‘fine’ inclusion of arterial segments and WSS*ref*= 5 Pa. 7B: adaptation dynamics for this network with WSS*ref*= 20 Pa. 7C: Initial (dashed lines) and steady state (solid lines) number of segments, sinks and loops (upper panel) and flow (lower panel) for this network for WSS*ref* between 1 and 30 Pa. 7D-E: Levels of WSS*ref* that lead to a reduction of the number of loops, segments, outflow connections (sinks) and flow to less than 50% of the initial values. Shown are results for nine mice from three strains, in ‘fine’ (7D) and ‘coarse’ (7E) networks.

### Mathematical analysis of loop stability

Of all the alternatives tested and in small networks as well as large coronary and cerebral arterial networks, no model based on adaptation to local hemodynamics resulted in stable loops in the simulations. This suggests that loop regression is a fundamental property of these classes of models.

Based on mathematical analysis, we indeed could prove that any topology for any set of the parameters and initial conditions regresses to a tree under WSS-dependent adaptation as defined by the model in eq. 1 (adaptation driven by WSS) or eq. 2 (generalized adaptation to a local hemodynamic stimulus). The proof is detailed in Supporting Information. In this analysis, we focus on the equilibrium points in networks with loops where all segments attain WSS*ref* and also have non-zero radius. We linearize the system around the equilibrium point by determining the Jacobian matrix and evaluate this matrix at the equilibrium. We then proof that the Jacobian matrix has as many eigenvalues with positive real parts as there are loops in the topology. This is the consequence of Kirchhoff’s’ circuit law, applied to pressure: in a closed loop, the directed sum of the pressure differences is zero. Using this we show that a linear combination of the variables of the linearized model increases as time increases. Hence the equilibrium of the linearized system is unstable and there exists a positive eigenvalue. The presence of an eigenvalue with positive real part for the linearized system implies that the equilibrium is unstable for the linearized system but also for the full nonlinear system [15].

We provide the proof for increasingly complex models: In the Supporting Information, we first analyze the model for WSS-driven adaptation (eq 1) for the triangular network (SI 2). We then extend this to the generalized adaptation model of eq. 2 for all *α*>*0* and *β*>0 in the triangular network (SI 3.1). Further mathematical analysis results in proof of loop instability for several models based on the WSS-driven model of eq. 1: direction-dependent adaptation (SI 3.2), heterogeneous WSS*ref* (SI 3.3), heterogeneous adaptation speed (SI 3.4). In SI 4 we show loop instability for the subsets of the general adaptation model under dynamic conditions, where one or more parameters oscillate around a mean value. We analyze this for oscillations that are either much faster (SI 4.1) or much slower (SI 4.2) than the adaptation dynamics. We show loop instability for all of these dynamic conditions. We finally show that loops are unstable in the generalized model applied to arbitrary network topologies that contain one or more loops (SI 5). Evidence for loop instability in combinations of these scenarios can be derived but is not shown.

## Discussion

### Summary of major results

Adaptation to wall shear stress is broadly considered to be dominant in the diameter regulation of arterial segments in a network. Such adaptation would lead to an equilibrium where WSS in all segments becomes equal to some reference value (WSS*ref*). However, the current simulation results and mathematical analyses show that this equilibrium, if it exists, is unstable in networks containing loops such as arcades and collaterals. Consequently, adaptation to WSS invariably leads to regression of segments in loops, transforming looping topologies into mere trees. This was found in simulations of a very wide variety of adaptation models, parameters values, initial states and oscillating parameters. Mathematical proof of loop instability was derived for most of these models. Table 3 summarizes these results.

**Table 3:** overview of the performed simulations and mathematical analyses. None of these revealed maintenance of arterial loops after adaptation.

|  | Adaptation to WSS | Generalized adaptation | Direction-dependent adaptation to WSS | Heterogeneous adaptation speed | Heterogeneous reference values for WSS | Dynamic conditions |
| --- | --- | --- | --- | --- | --- | --- |
| Simulation in small networks | & | & | & | & | & | & |
| Simulation in coronary and cerebral networks | & | Not tested | Not tested | Not tested | Not tested | Not tested |
| Mathematical proof for loop instability (general networks) | & | & | & | & | & | Proof for specific conditions |

### Modeling choices

We started with two very simple topologies, the triangle and cross-connected pentagon, allowing the study of a large array of adaptation models, with large variations of initial states and parameter values. After establishing the loss of loops here, adaptation was simulated in large coronary and cerebral artery networks that were based on experimental data. For practical reasons, this was limited to control of only WSS rather than the generalized model, and parameter variation was limited to WSS*ref*. Yet, based on the mathematical analysis, we are confident that all adaptation models and parameter sets would have resulted in regression of loops in these large networks.

Obviously, there are endless variations in adaptation rules, and we cannot rule out that there exists an adaptation model to local hemodynamics that results in stable loops. However, the mathematical analysis reveals that loss of loops is a fundamental consequence of the dependency between the node pressures in a loop, i.e. Kirchhoff’s circuit law. It seems that including more realistic models, such as inclusion of non-Newtonian properties of blood [16], will therefore not lead to fundamentally different results.

We did not consider development of new segments that could restore the loops, leading to a dynamic topology where loopiness is maintained by a balance between angiogenesis and regression. Such changes in topology are seen in the smallest vessels during development, in inflammation and in tumor vascularization. However, topology seems rather stable in healthy fully developed arterial networks, and a dynamic equilibrium of regressing segments and new connections restoring loopiness seems unlikely in at least the larger vessels.

We assumed that all segments in the networks similarly adapt their diameters, i.e. that none of the segments is a ‘special segment’ that closes the loop and that has different adaptation mechanisms. This seems reasonable for arcading networks such as the Circle of Willis and the mesenteric arcades (Fig.1), where all segments have comparable diameters and where it seems arbitrary which of the segments closes the loop. If these segments are indeed equal in terms of diameter regulation, it follows that none of these segments is controlled by WSS or other local hemodynamic conditions exclusively. In contrast to these arcades, pre-existing collateral segments, such as in the coronary circulation, are much smaller than the main branches that they connect. We cannot rule out that such collaterals regulate their diameter by other mechanisms, with the major coronaries still being regulated by exclusively WSS.

### Alternative mechanisms of diameter regulation

On average, arterial WSS is fairly similar over a large range of branching orders, covering a ∼100-1000-fold diameter range [9]. Yet, the observed variation between individual segments may well reflect the influence of other factors in regulation of vascular caliber[17]. Below we discuss these factors and their possible role in obtaining loop stability.

While the adaptation models depend on pressure gradients, we did not consider local pressure (P) itself as a stimulus. An acute increase in pressure causes vasoconstriction in seconds, a process known as the myogenic response [18]. Chronic hypertension is related to inward remodeling of predominantly the resistance arteries [19]. In both responses, wall stress (*σ*) is a possible stimulus. According to the Laplace relation, *σ* = *P*. *r*⁄*w* with *r* the radius and *w* the wall thickness. Accounting for wall stress as (additional) stimulus would require a more extensive model that includes more state variables, including wall thickness. We previously developed a 5-state adaptation model that covers biomechanics, wall thickness, wall composition and adaptation to both wall stress and WSS [20]. In initial work (in preparation) we applied this to honeycomb-like topologies and again invariably found loop regression. However, future work should address this in more detail.

Blood flow and flow reserve are matched to the metabolic needs of the tissue. Such autoregulation involves local tissue-derived stimuli. Effects of such metabolic stimuli as extra input to the diameter control was implicitly covered by considering a range of reference values for WSS, as well as by including heterogeneous WSS*ref*. However, we did not include full metabolic feedback loops here that aim to control oxygen delivery to the downstream tissue. Future work should address this. For quite some networks (circle of Willis, the mesenteric arcades) the segments are not in direct contact with the perfused tissue, making this metabolic influence unlikely.

Segments in arterial networks are known to communicate through electrical coupling or second messenger signaling between endothelial cells or other cells in the vascular wall [21]. Such communication is now considered important in acute regulation of vasomotor tone in the neurovascular unit, where signaling from neurons and astrocytes to capillaries results in vasoactive responses in upstream vessels [22]. While this concerns acute regulation in seconds, we [23, 24] and others [25] have provided compelling experimental evidence that long-lasting levels of vasomotor tone drive inward and outward remodeling processes. It seems possible that spatiotemporal integration of responses to local (hemodynamic) stimuli provides loop stability. These processes were considered by Pries et al. in what is known as the ‘shunt problem’. Here, short connections between arterial entrance and venous exit vessels are at risk of entering a vicious circle of increased flow, higher WSS and outward remodeling [26, 27]. In modeling work, these authors found that avoiding such shunting requires information transfer through coupling between segments. Coupling needed to be unidirectional, upstream in the arteries and not ‘around the corner’ in a node, i.e. from upstream in one segment to downstream in its sister. There is no evidence for unidirectional signaling capacity in acute regulation. Unidirectional signaling in structural remodeling might possibly relate to migration of endothelial cells against the flow, as observed *in vitro* and during embryonic development [28]. Edgar et al. [29] simulated such migration using cellular automata. In these models, migrating endothelial cells need to decide which path to follow in a flow-converging (‘venous’) node. They observed loop stability for decision rules that invoked both flow and endothelial cell numbers in the two legs. A concern here is the relevance for mature arterial networks, where caliber depends on the organization of the elastin and collagen in the wall rather than the number of endothelial cells. Clearly, the existence and nature of unidirectional coupling warrants future experimental work, notably because differences herein may exist in disease states [17].

### Implications

Individual differences in collateralization and completeness of arcades are substantial and cause large differences in outcome in acute ischemic stroke [30] and myocardial infarction [31]. While clues for genetic causes of these differences have been identified in rodents [32], this is not the case for humans. A better understanding of the mechanisms that lead to maintenance of loops could provide diagnostic tools and risk stratification in patients. We believe that this requires experimental work but also a network analysis. The current work aims to contribute to this.

Several arterial network models have been developed, frequently for the prediction of tissue perfusion [33, 34] and interpretation of CT perfusion and MR perfusion imaging [35]. While actual data can be used over a limited length scale, full arterial network models require application of branching rules or design principles [36]. The current work argues against regulation of only local WSS as a design principle. A better mechanistic understanding of adaptation may lead to more realistic algorithms for network generation. Such algorithms are becoming more and more relevant for setting up digital twins and in silico trials [37], while unraveling mechanisms of network adaptation is also of increasing relevance for developing perfused artificial organs [38].

### Limitations

This is a theoretical study. We covered a very large range of adaptation models and provided ample simulation results and mathematical prove for instability of arterial loops. However, many alternative models are possible. We do believe that the current work provides sufficient ground for warranting experimental work on the mechanisms that regulate arterial caliber in networks, with emphasis on direct (electrical) communication between segments in a network.

## Conclusion

The current computational and mathematical analysis reveals a fundamental issue in arterial network adaptation: while WSS and related hemodynamic stimuli are commonly believed to be the main drive for regulation of vascular caliber, loops such as arterial arcades and collaterals are not stable under such adaptation. Since these structures clearly exist, a search for the underlying mechanisms of loop stability is warranted.

## Methods

### Definitions

Arterial networks were generated consisting of sources, nodes and segments (Fig. 2). Sources connect to the outside world, carrying inflow or outflow. These were defined as combinations of source pressure and source conductance and are not adapting. Segments connect two nodes and are the adapting elements in the models. Nodes connect one or more segments or sources. Arterial loops were defined as areas fully enclosed but not intersected by segments. In the right example of Fig. 3 there are two loops, spanned by respectively segments 1-2-6 and 3-4-5-6; the area enclosed by segments 1-2-3-4-5 is intersected by segment 6 and is not considered as a separate loop.

### Arterial topological models

Vascular adaptation and network stability were analyzed for a series of arterial network topologies. These networks include artificial small loop models (3 or 6 segments) and extensive networks based on observed human coronary and mouse cerebral data, containing in the order of 10^4^ – 10^5^ segments.

#### Small loop models

The small topologies of Figs. 2-3 were used for extensive analysis of network adaptation for a range of initial conditions, parameter values and adaptation rules.

#### Human coronary arterial network

A further arterial network model was based on the human coronary arterial network. These data were taken from Schwarz et al. [11]. In that study, the left coronary arteries of a 84-old female who had not suffered from major cardiovascular events were filled with fluorescing casting material, followed by 3D imaging using an imaging cryomicrotome [12], segmentation, skeletonization and automated segmental diameter estimation. This network includes ∼ 200,000 segments with radii between 2 mm and 15 µm. The authors also established estimates for the ∼ 100,000 sink conductances representing the distal arterioles and downstream capillaries and veins. We included this full network in our adaptation model. The authors further estimated hemodynamics and WSS, based on the non-newtonian properties of blood (Fåhraeus-Lindqvist effect). For segments with radii of 15 µm and higher, the effective viscosity at a normal hematocrit varies by ∼25% over the diameter range [39]. We ignored this in order to limit the extensive simulation time, taking blood as Newtonian with a viscosity of 0.0035 Ns/m^2^.

#### Mouse cerebral arterial networks

We used data provided in Paetzold et al. [14], based on [13], which consist of nine mice (3 BALBc mice, 3 C57BL/6 mice, 3 CD1-E mice). These data consist of graph representations of the arterial and capillary network connectivity and radii. The original datasets contained around 5 million segments for each mouse brain. The vast majority of these segments represent the capillary bed. We selected arterial networks from these data as follows. Firstly, all segments with radius below a threshold value of 30 or 60 μm were removed. This resulted in several large networks, originating from the Circle of Willis, as well as in many small networks with radii around these thresholds. These small networks had become disconnected, e.g. due to removing the middle of three consecutive segments with radii of 17 – 14 – 16 μm for the 15 μm threshold. Therefore, secondly, we reconnected these isolated networks by adding again the 14 μm segment. This was done for all isolated networks with at least two segments. While in this example the procedure is straightforward, a more generalized reconnection procedure was needed due to the complex topology. For this we used the ‘shortestpath’ function in the matlab graph toolbox, using the conductances of the segments as weight. The reconnection procedure resulted in the inclusion of very many segments that had estimated radii below the chosen thresholds. We label the networks as ‘fine’ or ‘coarse’ for initial cut-off radii of respectively 30 and 60 μm.

Each pruned connection to the capillary network was replaced by a single non-adapting sink conductance. The conductances of these sinks, *G*_*s,i*_, were all taken the same:

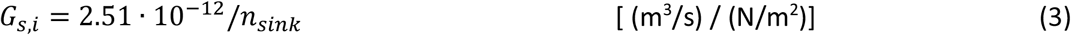

where *n*_*sink*_ denotes the number of such sink connections. This value is based on estimates of total flow (2 ml/min) and pressure gradient (100 mmHg) in the mouse cerebral circulation.

### Arterial hemodynamics and adaptation model

#### Physical relationships and adaptation models

Here we use double indices for the adapting segments, with segment *jk* connecting the nodes *j* and *k*.

Assuming steady perfusion by a Newtonian fluid, conductance of individual segments was calculated based on Poiseuille:

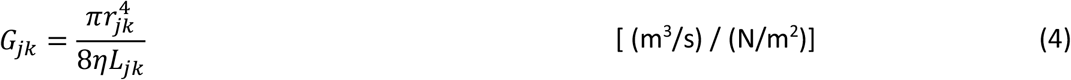

where *G*_*jk*_, *r*_*jk*_ and *L*_*jk*_ are the conductance, radius and length of segment *jk*, respectively. *η* is the fluid viscosity. In order to derive the nodal pressures from the conductances and source pressures, Kirchhoff’s current law, equivalent to conservation of volume, was applied for each node *j* in the network:

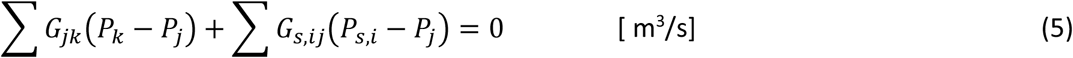

*G*_*s,ij*_ is the conductance of source *i* connected to node *j. P*_*k*_ is the pressure at node *k. P*_*s,i*_ is the source pressure in source element i. The summations are over all existing connections, where each term reflects a flow towards node *j*. This set of linear equations was solved for all *P*_*j*_ by matrix inversion.

Wall shear stress (τ_*jk*_) in each segment *jk* was derived from the pressure difference, radius and length:

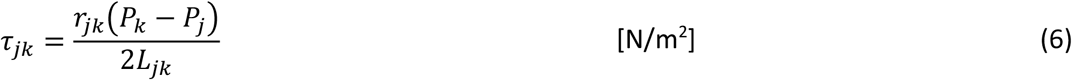

Biological adaptation models were defined in which the segmental radii change over time in response to the difference between a local hemodynamic stimulus and its assumed reference value. For adaptation to local WSS, this model is given by

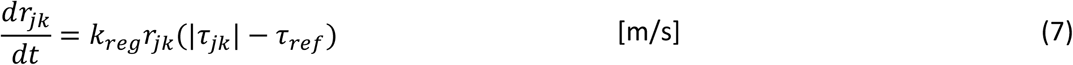

That is, individual segments become wider if their absolute WSS exceeds the reference value, τ_*ref*_, and shrink if their WSS is below the reference. We take the absolute value of WSS since the direction of positive flow is arbitrary and since we are not aware of experimental evidence that the response to maintained WSS depends on the direction. Note that, while the adaptation model is based on a local stimulus only, local WSS is a network property, leading to interaction of adaptation between segments.

We further considered a generalization of the adaptation model by introducing a stimulus *x*_*jk*_ that depends on the radius and pressure gradient as

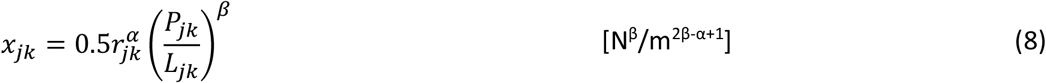

With α and β >0. The adaptation model then generalizes to

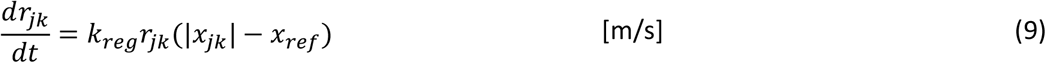

Specific models are α=2, β=1, where the stimulus is proportional to velocity. For α=4, β=1, the stimulus is proportional to flow. For α=4, β=2, the stimulus is proportional to power dissipation per vessel length. The proportionality constants involve the viscosity, which is kept constant in the simulations.

#### Application to the small loop models: dimensionless parameters and parameter space reduction

For application to the small loop topologies, the above model was converted to a dimensionless model with less parameters as follows:

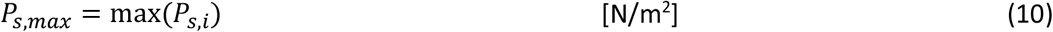

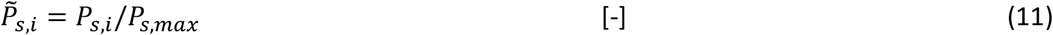

Where *P*_*s,i*_ is the source pressure of the i^th^ source element, and the maximum is taken over all source elements. 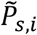 is the pressure at source *i* rescaled to the maximum. We assumed min(*P*_*s,i*_) to be zero; the current models only involve pressure gradients and no absolute pressures, rendering them invariant under a simultaneous shift of all source pressures.

We rescaled the conductances as

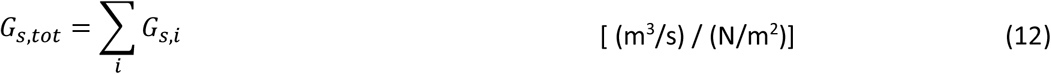

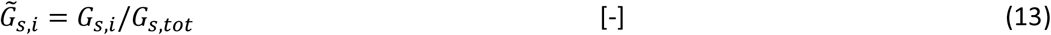

Where *G*_*s,i*_ is the conductance of the i^th^ source element, and the summation is over the conductances of all source elements. 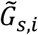 represents conductance rescaled to the sum of source conductances.

The length of the segments is rescaled by

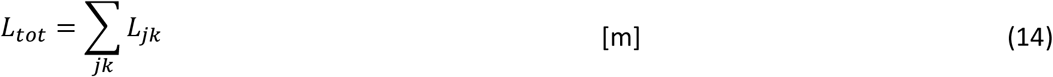

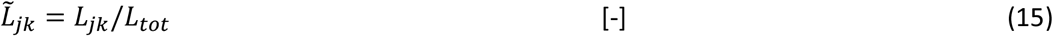

Where *L*_*jk*_ is the length of adapting segment *jk* in the model, connecting nodes *j* and *k*, and the summation is over all adapting segments. 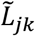 is the segment length rescaled to the total length of all segments.

We rescale the radius by

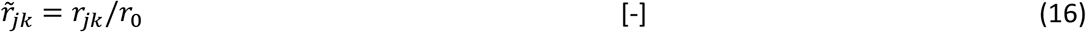

With *r*_0_ a typical radius in [m] set for scaling, and 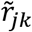 the dimensionless rescaled radius of segment *jk*.

We rescale the wall shear stress and general hemodynamic drive by

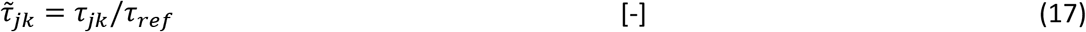

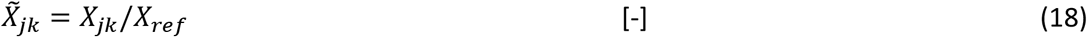

where 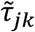 and 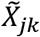 are the WSS and general drive, rescaled to their reference values.

A number of parameters can be lumped into three parameters:

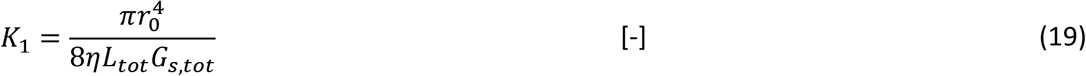

*K*_1_ reflects the balance between source conductances and network conductances.

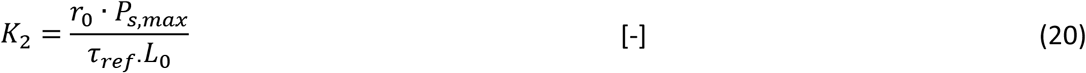

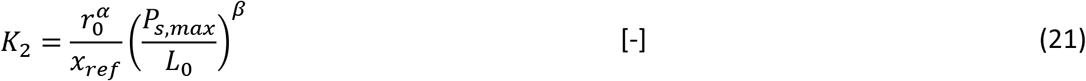

*K*_2_ indicates the balance of the pressure drive and reference for adaptation, indicated for respectively the WSS model and the general adaptation model.

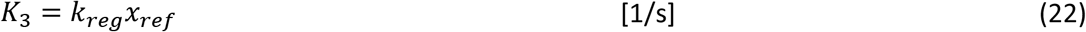

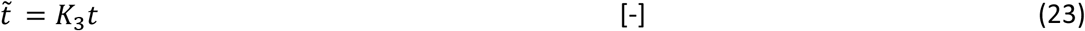

*K*_3_ defines the dynamics of adaptation. Different values of *K*_3_ cause a mere scaling of the dynamics over time; 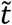 is the dimensionless time rescaled to these dynamics.

With this rescaling, the original hemodynamic relations and adaptation model become:

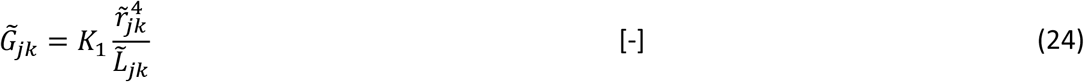

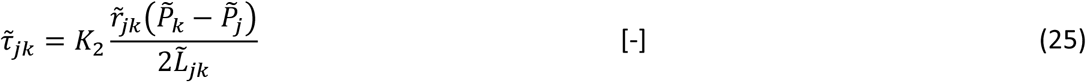

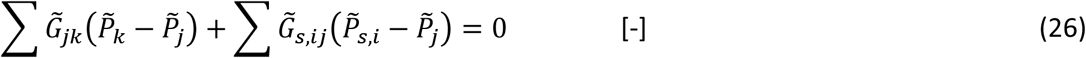

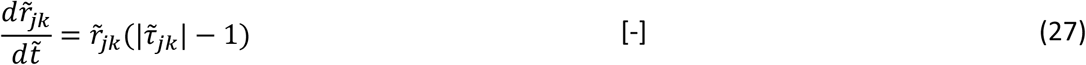

And for the general case:

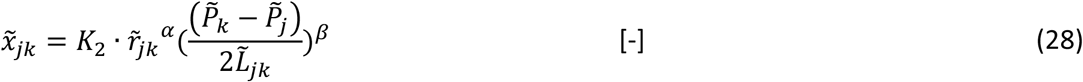

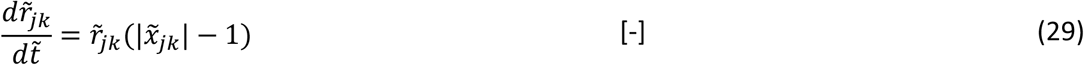

The triangular network example requires 7 dimensionless parameters: 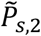 for one source (since the other two are 1 and 0 by definition), two values for 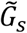 as well as for 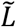 (since the rescaled values for the three sources respectively the three lengths both count up to unity), *K*_1_ and *K*_2_. The model predicts the state (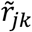 of all segments) and derived variables, including 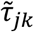, as a function of 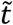

#### Estimation of parameter ranges for the small loop models

Default values for the various parameters (Table S1) and ranges for parameter variation (Table S2) were taken as follows:

The rescaled source pressures, 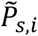, cover the range [0,1] by definition. We took equal intervals between the individual sources as default choice, e.g., 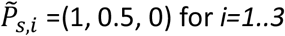 in the model with three sources, with variation of the middle source in the range [0.05,0.95] for the triangle model and [0.25,0.75] for the pentagon.

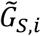 and 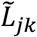 are in the range [0,1] by definition. As default model parameters, all 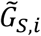 were taken equal 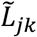 were taken unequal to bring in asymmetry in the models. Parameter variation included choices for specific sets of these values, covering the range [0.05,0.95] for the triangular topology and [0.25,0.75] for the pentagon.

For *K*_1_ we considered that the models form a part of a larger network and therefore the default model should have a balance between (initial) conductances of sources and segments. For higher *K*_1_ the source conductances, relative to the segment conductances, are lower (eq. 19), and become more and more dominant in determining the network pressures and flows. This balance depends in a complex way on the network topology. As a ballpark, we took *K*_1_ = 1 as the default value. This represents a model where neither the source conductances nor the (initial) network conductances are dominant in determining local pressures and flows. In the models, we covered the range [0.1,10] for *K*_1_.

For *K*_2_ in the WSS model (eq. 20 and 21) we considered *r*_0_ = 10^−4^ m, reflecting the larger resistance vessels (36), with their lengths being 20 times the radius, or 2·10^−3^ m. This could reflect a loop of rat mesenteric resistance arteries. Observed WSS values in such arteries are in the order of 5 N/m^2^. With pressure gradients over these networks in the order of 10% of the systemic pressure of 13,000 N/m^2^, *K*_2_ would be 13. We took *K*_2_= 10 and 20 for respectively the triangle and pentagon topologies as default and covered a 100-fold range for *K*_2_ as parameter variation. For the general adaptation rule, *K*_2_ = 20 was used as default for both topologies.

By expressing all results as function of a rescaled time 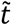, there was no need to choose *K*_3_. The processes considered here are based on vascular remodeling by WSS, a process that typically occurs in the time domain of weeks.

### Simulation of small network adaptation

Simulations were based on the hemodynamic relationships and adaptation laws depicted above and were implemented in Matlab R2022b. These were performed using a 4^th^-5^th^ order Runga Kutta method with a reduction in maximum integration step size where needed and otherwise default settings. For adaptation to WSS (eq. 1) in the small loop models, initial rescaled radii for these simulations were based on derivative-free searches. In these, we minimized the difference between the WSS and its reference value over all segments. More specifically, we chose the initial radii based on the sums of squares of deviations of the rescaled WSS from its reference of 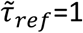 :

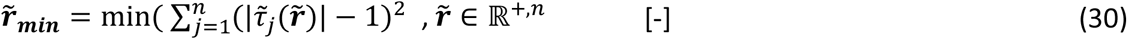

The summation is over all 3 or 6 adapting segments *j* in the small networks (see insets in Fig. 3 for numbering). 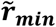 is the set of radii providing the minimum deviation. The search was done by the ‘fminsearch’ function in Matlab, which uses the Nelder-Mead simplex algorithm. The search started at all 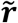 equal to unity. For most parameter settings, the minimum was equivalent to 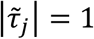 in all segments, denoting a (possibly unstable) equilibrium state. A similar procedure was followed for the general model (eq. 2)

In default, initial radii for the simulations were set to 1% above 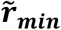. In order to study the effect of initial conditions, initial radii were typically varied in the range [0.3,3] times 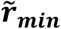, with a more extensive variation of [0.05,4] times 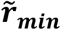 for the default triangular topology. Dimensionless simulation time was typically 30, long enough to include development of a steady state as judged from the final deviation of 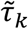 from unity in remaining segments. Simulations with slow oscillations were continued long enough to cover several periods. Sets of simulations included sweeps of initial radii or model parameters.

### Simulation of coronary and cerebral arterial network adaptation

For simulation of the large networks, actual rather than dimensionless parameters and variables were used. Simulations were performed based only on the WSS adaptation model of eq. 1, with variation of WSS*ref* between 1-30 N/m^2^. In the coronary artery network, initial radii were set to the published values. In the cerebral networks, as indicated by the authors [14], the estimations of radii are insufficiently reliably for a hemodynamic analysis. We therefore set all initial radii to an arbitrary value of 70 μm. For both networks, we assumed that adaptation occurs in the course of months and set *k*_*reg*_ = 0.03 Pa^-1^day^-1^ = 3.45·10^−7^ Pa^-1^s^-1^. Source pressure was taken as 13.3·10^3^ Pa=100 mmHg. Simulations were performed by using the 4^th^-5^th^ order Runga Kutta method. We simulated an adaptation period of 333 days.

We analyzed changes in topology and hemodynamics during network adaptation, as well as the steady state numbers of loops, segments, connections to the downstream vessels and flow. The number of loops in these networks (nL) was calculated from the number of segments (nS) and nodes (nN, including the source and sink nodes) according to

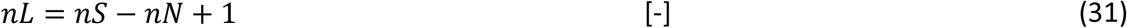

This relation holds for a fully connected network, and is based on Euler’s Polyhedron formula on the number of faces.

Simulations of these large networks was done on a Windows computer with an Intel Xeon E5-2630V4 2.20 GHz CPU, requiring several hours per network on a single core. Simulations of different networks were done in parallel, using up to 12 cores.

### Mathematical analysis of arterial loop stability

The supplement provides the mathematical stability analysis of equilibrium points for adaptation.

## SUPPORTING INFORMATION

**Table S1:**
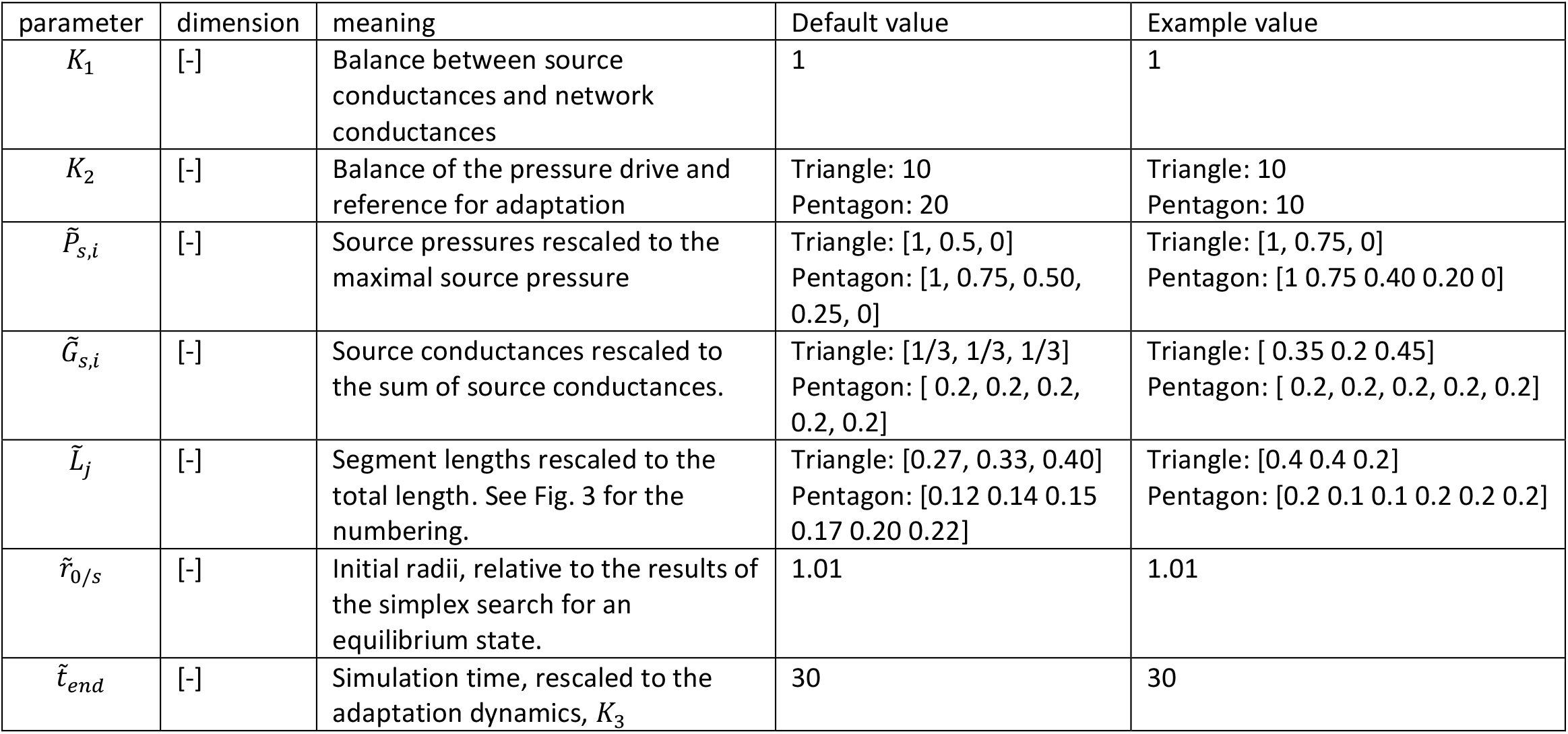
Small network model parameters for the default simulations and for the examples of Fig. 3.

**Table S2:**
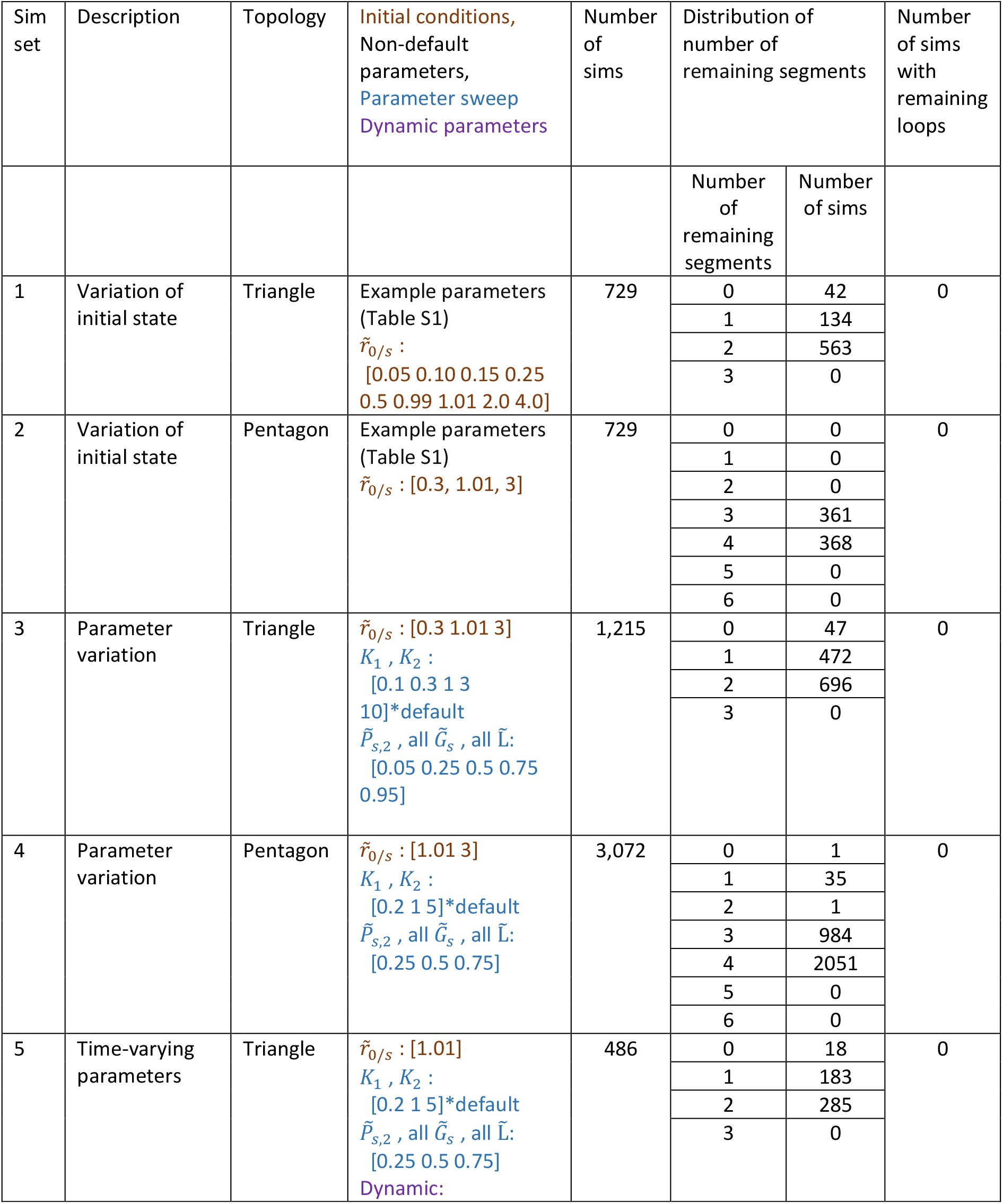
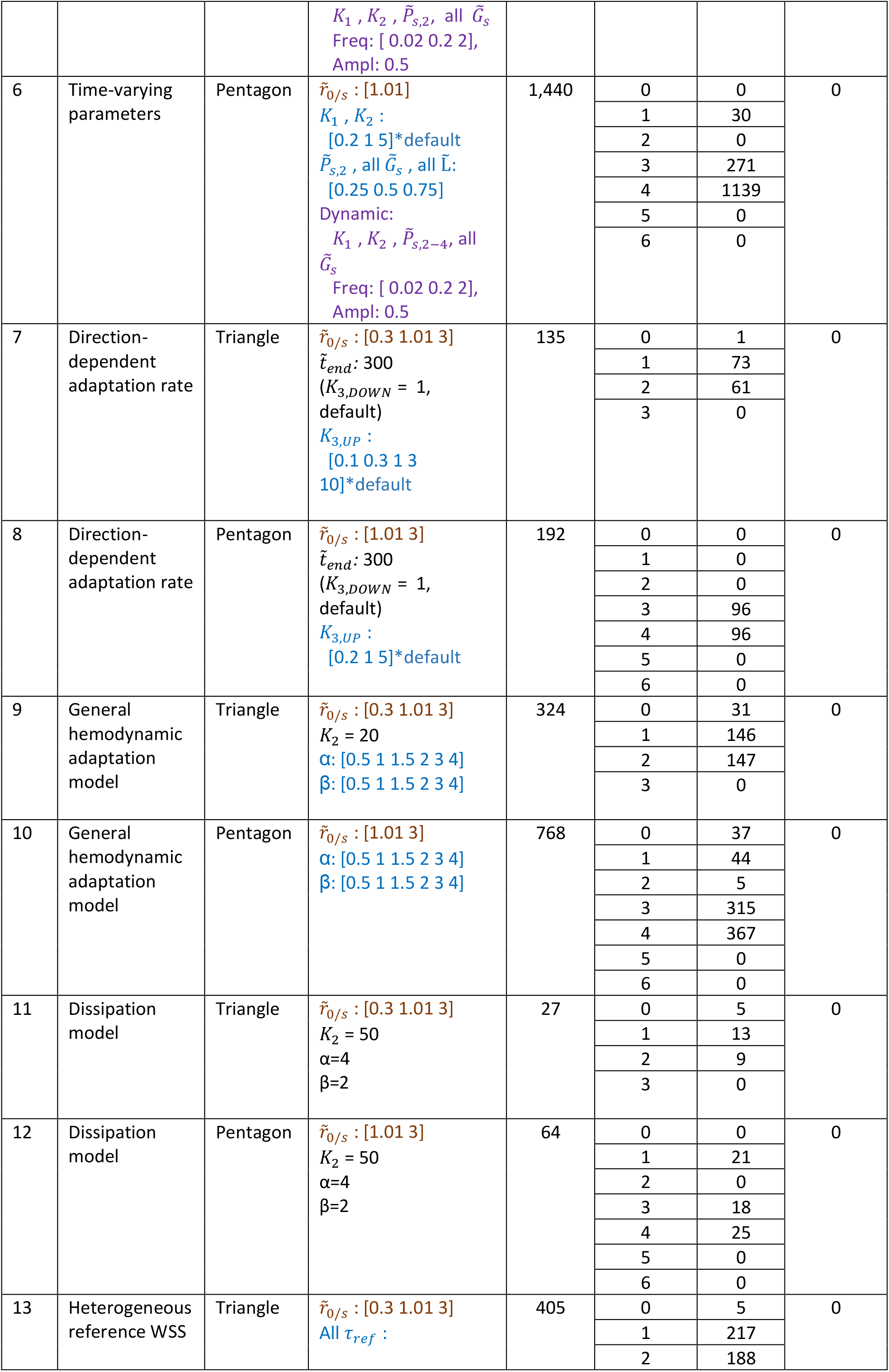
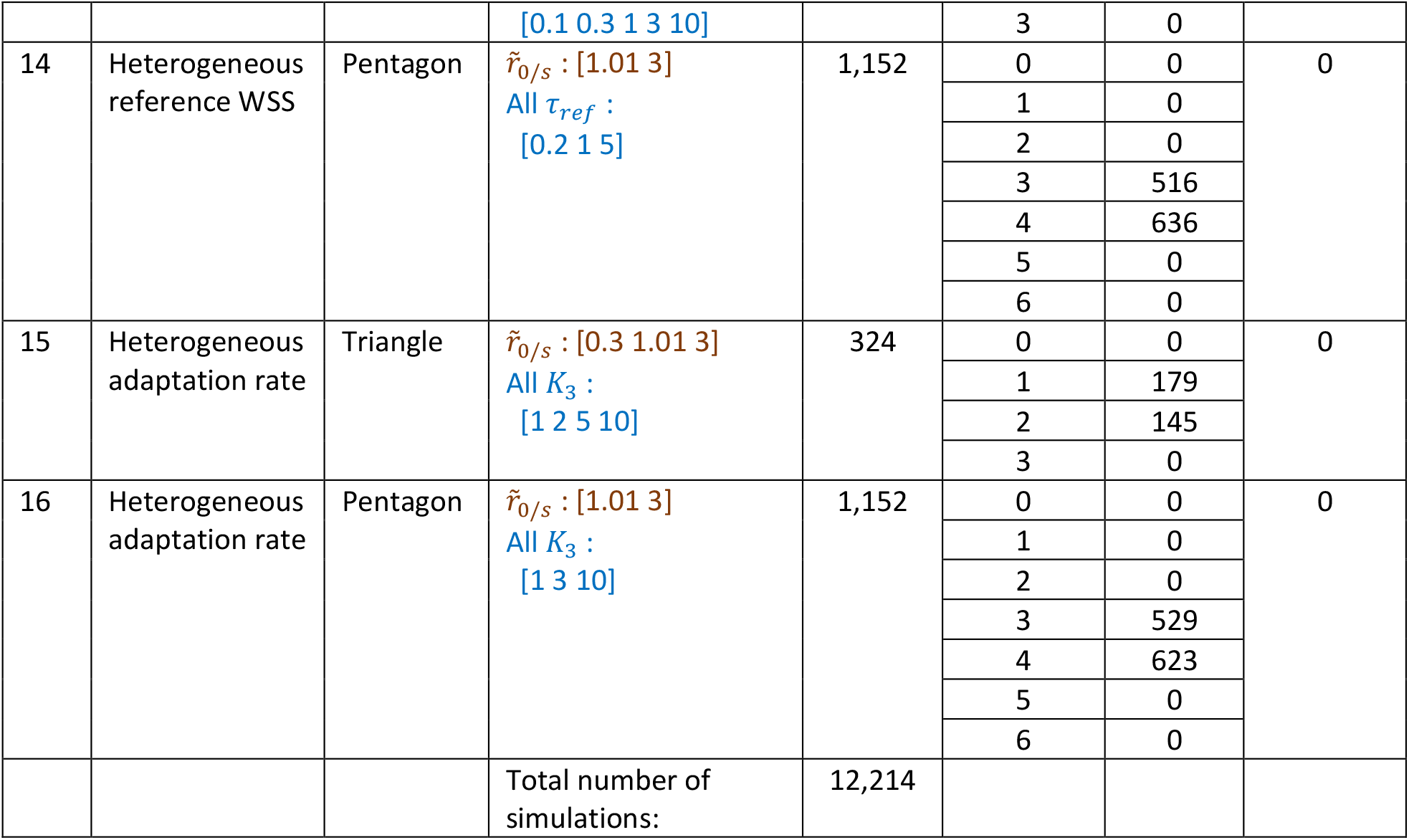
Simulation results. Parameters are at default values unless otherwise indicated. 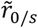 : initial relative deviations of radii from results of the simplex search for an equilibrium. All indicated combinations of 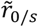 for the adapting segments were simulated, e.g in simulation set 3, for 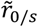 : *[0*.*3, 1*.*01, 3]*, 27 combinations of initial states were simulated. This set of initial conditions was used for each variation in the other parameters. Parameter sweeps were done by varying each of the indicated parameters while keeping the others at their default value. For the dynamic parameters, the choices for initial states and several time-averaged values for parameters were combined with a range of sine-wave oscillations of amplitude 0.5. For the direction-dependent adaptation rates, we changed the values for the rate of outward growth (*K*_3,*UP*_), while keeping the rate of inward adaptation or shrinkage at the default value of *K*_3,*DOWN*_ = 1. The results are summarized as the number of simulations that rendered a specific number of remaining segments, and as the number of remaining loops observed in these simulations.

Supporting movie Coronary adaptation: Regression of coronary segments for increasing levels of WSS*ref*. Distal conductance in these simulations was taken equal to the published values. Shown are steady state topologies for increasing WSS*ref* in each frame. For clarity, segmental diameters are not shown. Panel A: 3D view. B-D: 1 cm transverse sections, taken at respectively 6.21-7.21 cm (B), 4.29-5.29 cm (C) and 2.37-3.37 cm (D) above the apex of the heart. Blue: open segments; red: segments that have regressed at the indicated WSS*ref*. Circles: entrance points (left anterior descending and circumflex arteries) of the left coronary circulation. The right coronary circulation was not included in the data.

## SI 1 Prove of the loss of loops in models for vascular adaptation

In this Section, we study the various models that were presented in the main text analytically. In these proofs we study the system presented in equation (2) in the main manuscript:

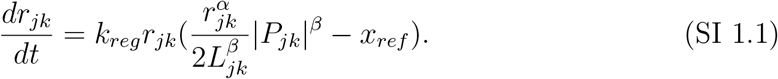

In this analysis the fact that this system is in the dimensionfull setting does not influence it; it does make the dependence on the parameters clearer. In order to analyse the stability of the equilibria, we need to differentiate |*P*_*jk*_|, thus, we need to be careful because of the absolute value. Therefore, we introduce the sign of *P*_*jk*_ as *s*_*jk*_ = sign (*P*_*jk*_).

## SI 2 Wall shear stress dependent vascular adaptation for the triangular topology

First, we assume adaptation to wall shear stress and take *α* = *β* = 1 and analyse the model for a triangular topology. The pentagon setting studied in the main manuscript can be treated in an analogous way, see also Section SI 5 for a very general network topology. The model for a triangular setting of vessels and *α* = *β* = 1 is given by

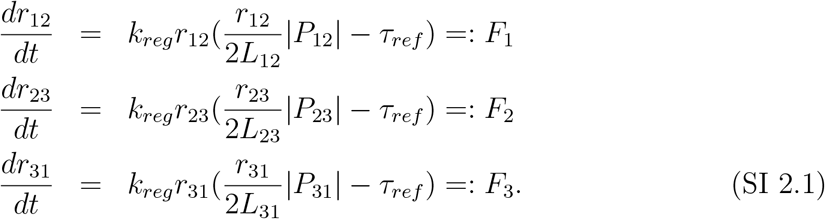

where *P*_12_ + *P*_23_ + *P*_31_ = 0.

Then, the fixed points need to satisfy

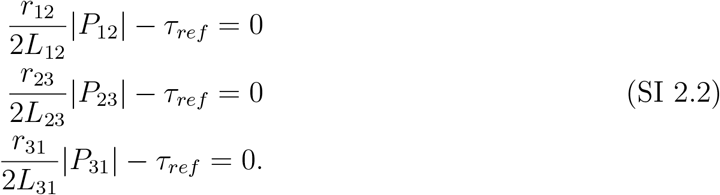

Now, we assume that the parameters are such that there exists at least 1 fixed point for which all *r*_*i*_ ≠ 0 and study that point from now onward. In case system (SI 2.1) does not have a fixed point with all *r*_*jk*_ ≠ 0 then we conclude that a loop is not present.

The stability of the fixed point can be determined by linearizing around the point, then we obtain

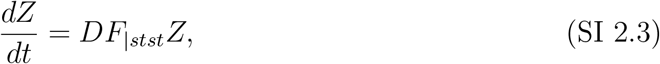

where *Z* = (*z*_12_, *z*_23_, *z*_31_) and *DF*_|*stst*_ is the Jacobian evaluated at the fixed point. And

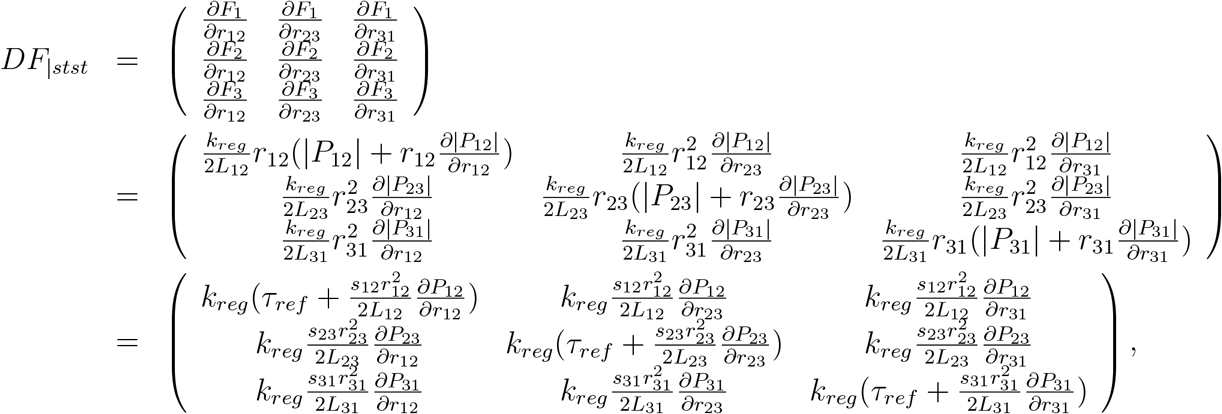

where the matrix is evaluated at the fixed point that satisfies (SI 2.2). In the last step we use that (SI 2.2) is satisfied. There are two methods to show that the fixed point is unstable.

### SI 2.1 First method

Using, equation (SI 2.3), we determine

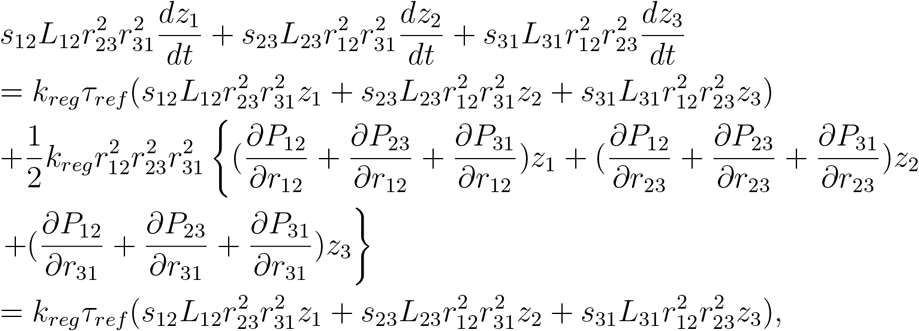

where the *r*_*i*_ satisfy (SI 2.2). In the last step we use that *P*_12_ + *P*_23_ + *P*_31_ = 0 such that differentiating this sum also yields zero. Next, we define

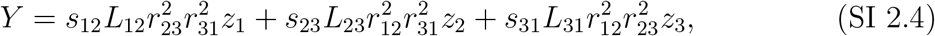

and obtain that

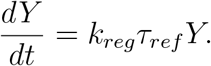

Solving this equation results in

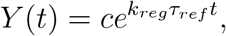

where *c* = *Y* (0) is a constant. Thus, for *t* > 0 and increasing *t* we find that *Y* also increases and we conclude that the combination

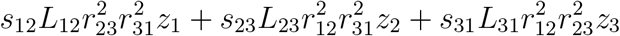

always increases. Moreover, we conclude that *λ* = *k*_*reg*_*τ*_*ref*_ is an eigenvalue and that it is positive. Therefore, we conclude that the fixed point *Z* = 0 in equation (SI 2.3) is unstable which corresponds to the fixed point (SI 2.2) being unstable for the linearized system. This also implies that the fixed point is unstable for the full nonlinear system (SI 2.1), see [2]. Hence, when a fixed point exists for which all *r*_*i*_ ≠ 0, it is unstable.

### SI 2.2 Second method

In this section, we will use another method to determine the stability of the fixed point (SI 2.2) namely by studying the eigenvalues of the matrix *DF*_|*stst*_. To obtain these eigenvalues, we need to determine the characteristic polynomial by analyzing

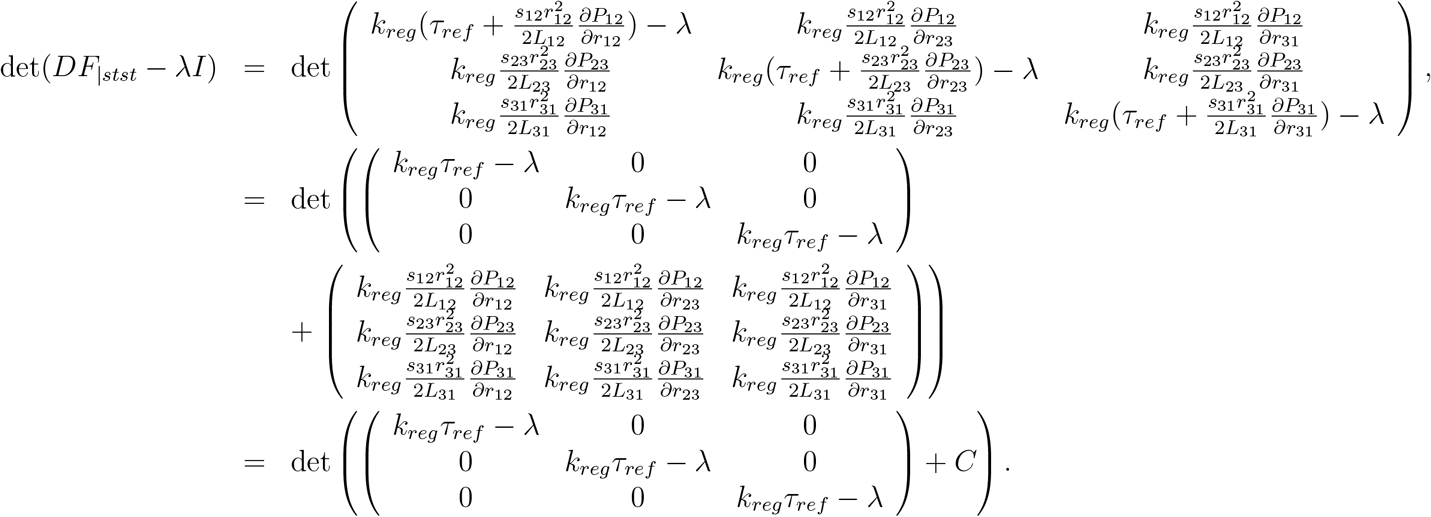

Here, *C* is a matrix of which the rows are linearly dependent since *P*_12_ + *P*_23_ + *P*_31_ = 0. We will use this fact to determine the above determinant.

#### Remark SI 2.1

*Note that in general it is not true that* det(*A* + *B*) = det(*A*) + det(*B*) *for 2 matrices A and B and so we do not use that here*.

If we write out the determinant we find that

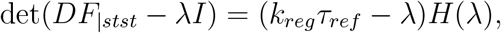

where *H* can be determined explicitly in terms of all above expressions. Next, we look at the solution of the characteristic polynomial

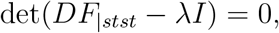

and conclude that one of the eigenvalues is given by *λ* = *k*_*reg*_*τ*_*ref*_ > 0. Therefore, one of the eigenvalues is positive, and thus, the fixed point is unstable also for the full nonlinear system (SI 2.1).

## SI 3 Loss of loopiness by local hemodynamic adaptation occurs in a wide variety of models

The above analysis can be used for a wide variety of models for hemodynamic adaptation which were also considered in the main text.

### SI 3.1 Generalized adaptation to hemodynamic conditions

In this section, we look at a generalized system for hemodynamic adaptation and still focus on the triangular setting. Thus, we analyse the system

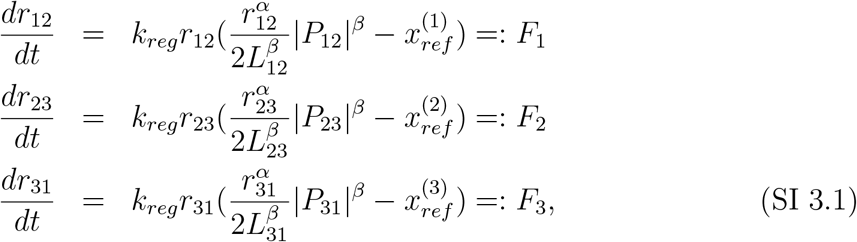

where *P*_12_ + *P*_23_ + *P*_31_ = 0, *α* > 0 and *β* > 0. Then, the fixed point for which all *r*_*i*_ ≠ 0 is given by the point that satisfies

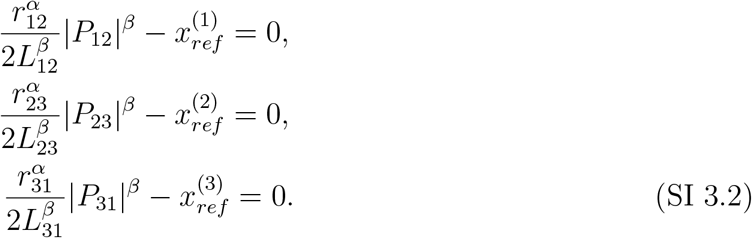

The stability of this point can be determined by linearizing around the point, then we obtain

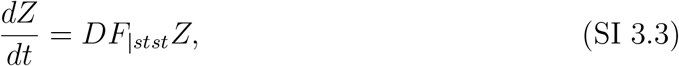

where *Z* = (*z*_12_, *z*_23_, *z*_31_) and

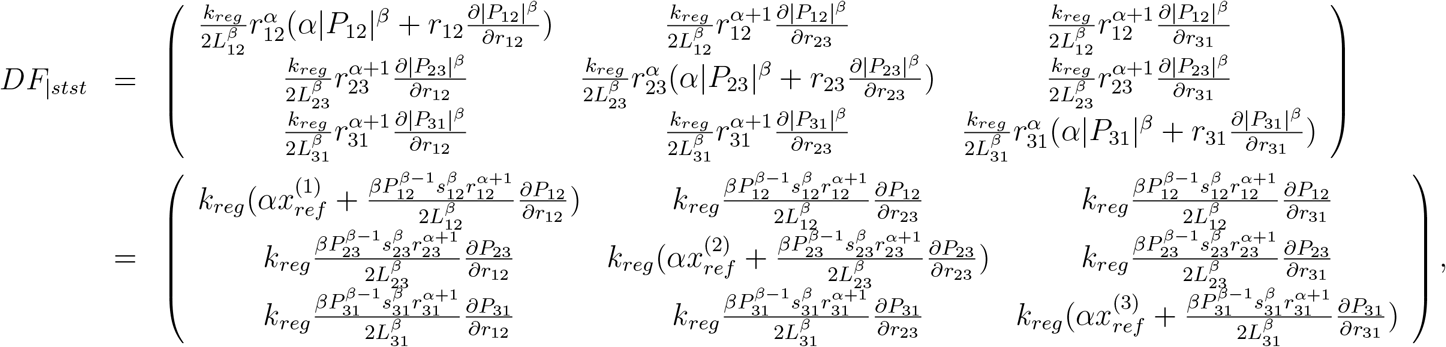

where the matrix is evaluated at the fixed point that satisfies (SI 3.2) and we again use this in the last step.

Here, we analyse the case where all the 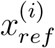 are the same so 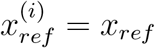 for *i* = 1, 2, 3. We use method 1 from Section SI 2.1 to prove that the fixed point is unstable. Method 2 from Section SI 2.2 could also be applied and results in the eigenvalue *λ* = *αk*_*reg*_*x*_*ref*_ > 0, we refrain from showing that here.

To apply method 1, we determine

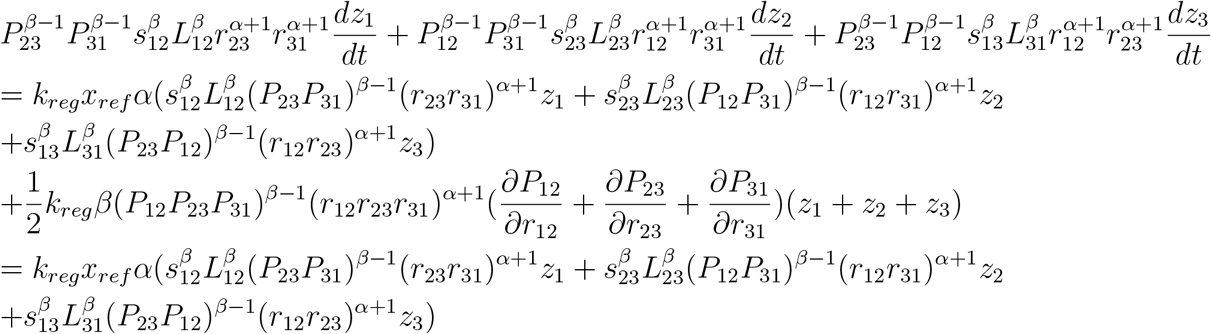

where the *r*_*jk*_ and *P*_*jk*_ are evaluated at the fixed point. In the last step we use that *P*_12_ + *P*_23_ + *P*_31_ = 0. Then, we find for

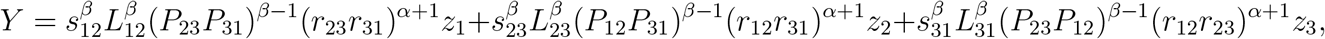

that

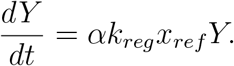

Solving this equation gives that

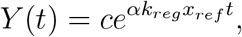

where *c* = *Y* (0) is a constant. Thus, for *t* > 0 and increasing we find that *Y* also increases and we conclude that the combination

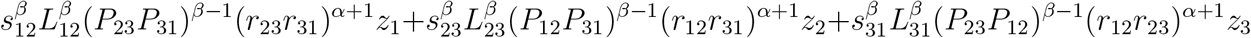

always increases. Similar to before this also results in the eigenvalue *α* = *αk*_*reg*_*x*_*ref*_. Therefore, we conclude that the fixed point *Z* = 0 in expression (SI 3 .3) is unstable corresponding to the fixed point (SI 3.2) being unstable for the linearized system and also for the full system (SI 3.1).

### SI 3.2 Different *τ*_*ref*_ among adapting segments

Here, we assume that the reference wall shear stress *τ*_*ref*_ are different for the various vessels and denote these by 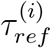. We can extend method 1 in section SI 2.1 to this case.

Analogous to before, we determine the linearized system and get that

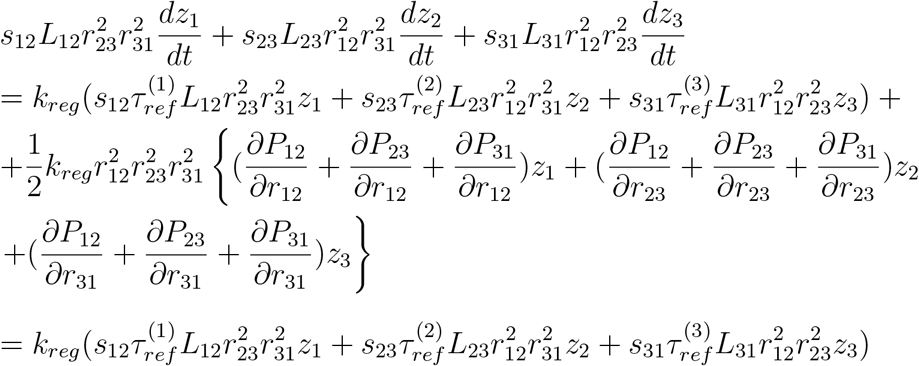

where the *r*_*i*_ are evaluated at the fixed point. In the last step we again use that *P*_12_ + *P*_23_ + *P*_31_ = 0.

Now, we assume without loss of generality that *s*_12_ = *s*_23_ = 1 and *s*_31_ = − 1 then there are three distinct possibilities:

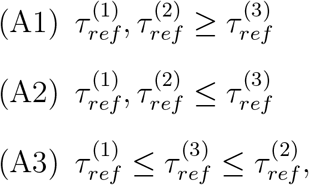

others are similar. We study case (A1) and assume, again without loss of generality, that 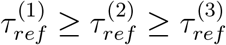

Then, we define

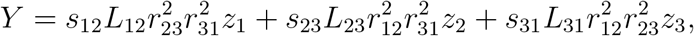

and find that

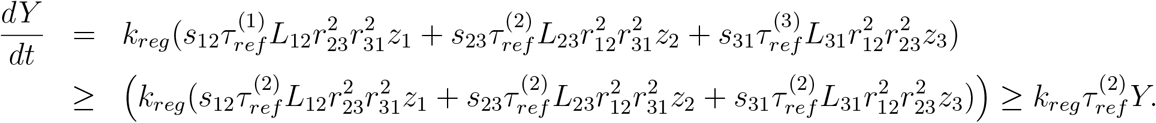

To determine an estimate for *Y* in this inequality, we can use a theorem by Petrovitch [3], see Appendix A for the precise statement. Applying the theorem, we conclude that

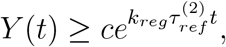

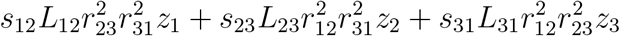

always increases. Therefore, we conclude that the fixed point *Z* = 0 in equation (SI 2.3) is unstable which corresponds to the fixed point (SI 2.2) being unstable, also for the full nonlinear system.

For possibility (A2), a proof analogous to the above proof can be given so we refrain for doing that here. When (A3) holds, we redefine the |*P*_*jk*_|’s in the system such that *s*_23_ = *s*_31_ = 1 and *s*_12_ = −1 and then a similar argument holds.

To summarise, the fixed point (SI 2.2) is always unstable in the case of different *τ*_*ref*_ ‘s.

### SI 3.3 Different adaptation speed between the adjacent segments

Here, we take a different adaptation speed for the segments, hence, the *k*_*reg*_ in the expressions for the *F*_*i*_ are taken different namely as 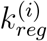. We can extend method 1 in section SI 2.1 to this case. Again, we determine the linearized system which is

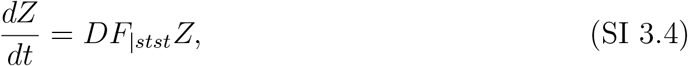

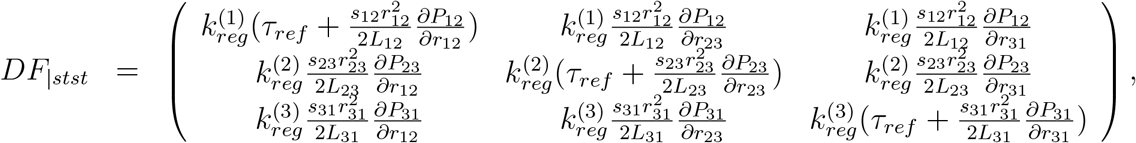

Using this, we determine

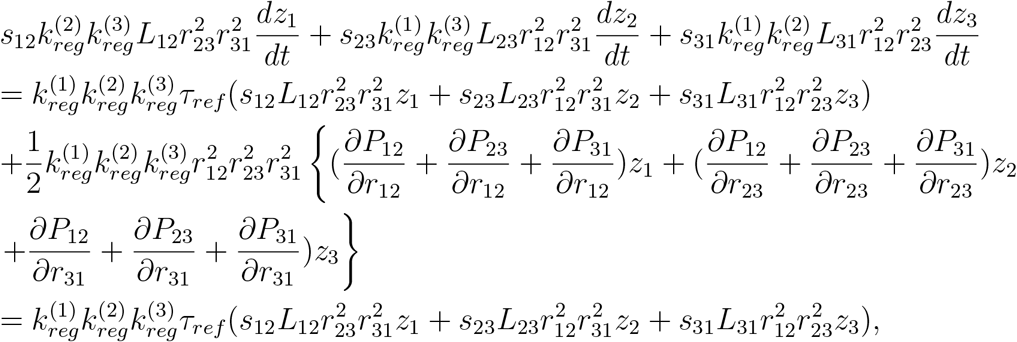

where the *r*_*i*_ satisfy (SI 2.2). In the last step we again use that *P*_12_ + *P*_23_ + *P*_31_ = 0.

Now, we assume without loss of generality that *s*_12_ = *s*_23_ = − 1 and *s*_31_ = − 1 then there are three distinct possibilities:

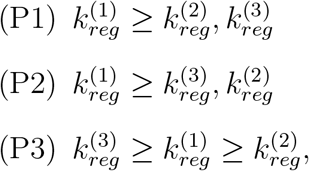

others are similar. We study case (P1) and assume, again without loss of generality, that 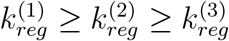

First, we study possibility (P3) then we find for

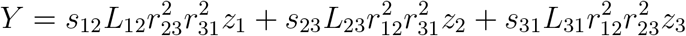

that

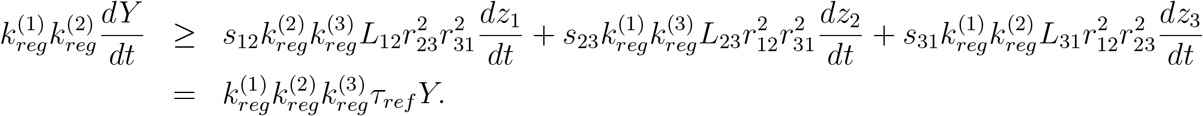

To determine an estimate for *Y* in this inequality, we use a theorem by Petrovitch [3], see Appendix A for the precise statement. Applying the theorem, we conclude that

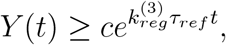

where *c* = *Y* (0) is a constant. Thus, for *t* > 0 and increasing *t* we find that *Y* also increases. Therefore, we conclude that the fixed point *Z* = 0 in equation (SI 2.3) is unstable which corresponds to the fixed point (SI 2.2) being unstable for the linearized system and also for the nonlinear problem. For the other possibilities we have to redefine the *P*_*jk*_’s again, similar to Section SI 3.2.

### SI 3.4 Direction dependent adaptation

Here, we analyse the case when the vessel radius increases slower and decreases faster than default. Hence

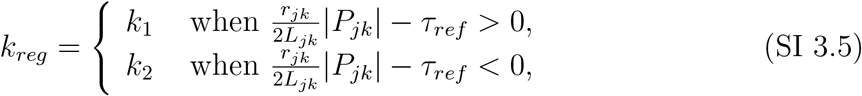

where 0 < *k*_1_ < *k*_2_ and we again choose *α* = *β* = 1.

In Section SI 3.3, we already showed that when the *k*_*reg*_ are all different (or when they are the same) that the equilibrium with all the *r*_*jk*_ ≠ 0 is unstable. Therefore, in that case, there exists at least one orbit which starts close to the fixed point but does not stay close to the fixed point for all time. More precisely, there exists an *ε* > 0 such that for every *δ* > 0 there exist initial conditions *r*_0_ = *r*(*t*_0_), *t*_0_ such that 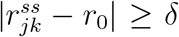 and 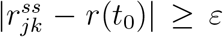 for all *t* ≥ *t*_0_ where 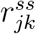 is the fixed point and *r*(*t*) is a solution of the system. Now, we assume that we start on this orbit. Then, the only possibility for the fixed point to become stable is when this orbit moves towards the fixed point. Hence, when it crosses one of the nullplanes 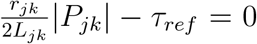 so that and the sign of 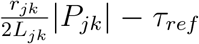 changes. However, that does not follow from the above assumption (SI so we conclude that the fixed point remains unstable.

## SI 4 Adaptation under dynamic conditions

Next, we analyse time dependent periodic variations of the parameters. We denote the vector of all parameters *G*_*s,i*_, *P*_*s,i*_ an *L*_*jk*_ by *µ*. When we take parameters to depend on time this implies that

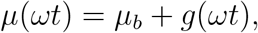

where *µ*_*b*_ is the vector of the constant input parameters and *g* is a periodic function with period *ωT*.

Moreover, we denote the vector of all radii *r*_*jk*_ by *R* and the corresponding system as

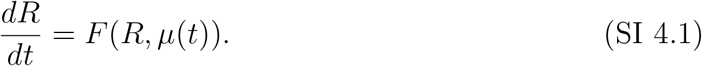

We analyse two choices namely when the periodic variation is slow and when it is fast. For the fast periodic variation we choose *ω* ≫ *k*_*reg*_*x*_*ref*_ whereas for the slow periodic variation we take *ω* ≪ *k*_*reg*_*x*_*ref*_.

Note that this system is the same as in the main manuscript since there we study the dimensionless system whereas we still analyse the system with dimensions.

### SI 4.1 Fast periodic variation of parameters

In the case when the perturbation is fast we assume that *ω* ≫ *k*_*reg*_*x*_*ref*_ and introduce 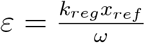 where 0 < *ε* ≪ 1. Then, we rescale system (SI 4.1) by introducing 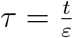 and obtain

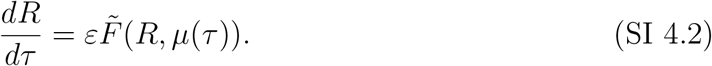

We will analyse this system by using the averaging method and introduce the associated averaged system

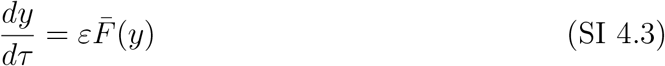

where

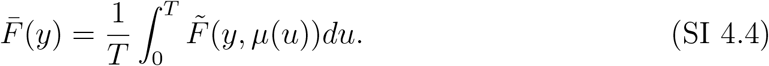

This system is of the same form as the system studied in Section SI 2, so with constant parameters. When we assume that there exists a fixed point (with all *r*_*jk*_ ≠ 0) then we already showed that for this system the fixed point is unstable.

To study the full perturbed system (SI 4.2), we can use the Averaging Theorem which can be found in for example [1], Theorem 4.1.1. Here we use that *F* is at least *C*^2^ and bounded on bounded sets, and of period *T* in *τ*. Then there exists a *C*^2^ change of coordinates *R* = *y* + *εw*(*y, τ*) under which (SI 4.2) becomes

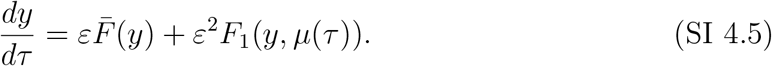

where *F*_1_ is of period *T* in *τ*. Moreover

i. if *R*(*t*) and *y*(*t*) are solutions of (SI 4.2) and (SI 4.5) with *R*(0) = *R*_0_, and *y*(0) = *y*_0_, respectively, and |*R*_0_ − *y*_0_| = 𝒪 (*ε*), then |*R*(*t*) − *y*(*t*)| = 𝒪 (*ε*) on a time scale 1*/ε*.
ii. If *p*_0_ is a hyperbolic fixed point of (SI 4.3) then there exists *ε*_0_ > 0 such that, for all 0 < *ε* < *ε*_0_, (SI 4.2) possesses a unique hyperbolic periodic orbit *γ*_*ε*_(*t*) = *p*_0_ + 𝒪 (*ε*) of the same stability type as *p*_0_.

Hence, we conclude from (ii) that in case the system (SI 4.5) has a fixed point with *r*_*jk*_ = 0 that system (SI 4.1) has a periodic solution for sufficiently small. Moreover, since the fixed point is unstable and hyperbolic (no eigenvalues with real part zero), the corresponding periodic solution is also unstable.

### SI 4.2 Slow periodic variation of parameters

In this section, we study the case when the parameters are perturbed by a periodic function that is slow, in other words, we choose *ω* ≪ *k*_*reg*_*x*_*ref*_. Then, the system is slaved to the original system.

We now set 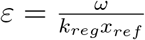 with 0 < *ε* ≪ 1. Then, by using a Taylor expansion, we know that there exists a function *g*_1_ such that

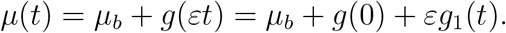

Substituting this in system (SI 4.1) and again using a Taylor expansion this reduces to

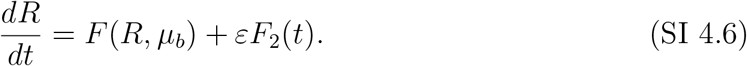

with *F*_2_ a function.

Now, for *ε* = 0 the system (SI 4.6) reduces to a system with constant parameters. Again we know from Section SI 2.1 that if there exists a fixed point with *r*_*jk*_ ≠ 0 it is unstable. Next, we use the Implicit Function Theorem in a similar way as in the proof of Lemma 4.5.1 in [1] to conclude that for *ε* sufficiently small, system (SI 4.6) has a periodic orbit. Moreover, we conclude that since the fixed point that exists for *ε* = 0 is unstable, the corresponding periodic orbit is also unstable.

## SI 5 General network that contains loops

The above can be generalised to a network that contains a loop. We assume that the network has n nodes, *K* in- or out-pressures denoted by *P*_*s,i*_, *i* = 1, ⋯ *K*, and contains and loop that consists of *m* nodes (*m* < *n*). Then, |*P*_*jk*_| = |*P*_*k*_ − *P*_*j*_| for two adjacent nodes and *k* ≠ *j*, otherwise (for non-adjacent nodes) |*P*_*jk*_| = 0. The system is still given by (SI 1.1) and here we study *α* = *β* = 1 but the analysis could also be extended to all the cases analyzed above. For the loop we use the notation *j*_1_, ⋯, *j*_*m*_ for the connecting nodes. So, *j*_1_ connects to *j*_2_ and so on, and *j*_*m*_ back to *j*_1_.

We find that the sum of all pressures differences within the loop is zero so

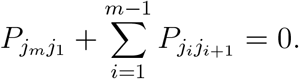

Again, we assume that the system (SI 1.1) has a fixed point were *r*_*jk*_ ≠ 0 for all adjacent nodes. Such fixed point exists depending on the choice of the parameters.

Linearising around the fixed point we obtain

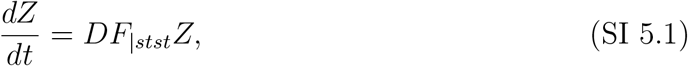

where *Z* is the vector over all nodes *z*_*jk*_ and *DF*_|*stst*_ is the Jacobian evaluated at the fixed point. Then, we define

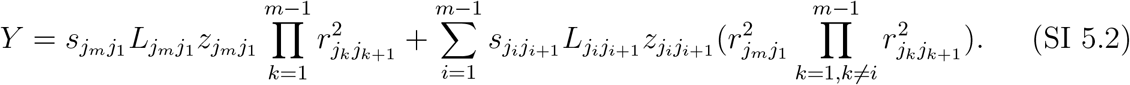

So we define an expression similar to (SI 2.4) but only take into account the vessels of the loop. Then, we find that

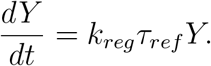

Solving this equation results in

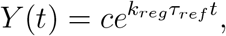

where *c* = *Y* (0) is a constant. Thus, for *t* > 0 and increasing *t* we find that *Y* also increases and we conclude that the sum (SI 5.2) over the *z*_*jk*_ in the loop increases. Therefore, we conclude that the fixed point *Z* = 0 is unstable which corresponds to the fixed point being unstable. Hence, the loop is unstable for the linearized and also for the full nonlinear system, and therefore, it will disappear.

### Remark SI 5.1

*In the case when there are more loops present in the network we can generalize the above and can conclude that all these loops are unstable and will disappear. This case relates to the eigenvalue λ* = *k*_*reg*_*τ*_*ref*_ *having a higher (algebraic) multiplicity and more eigenvectors so a higher geometric multiplicity*.

### Remark SI 5.2

*The above proof for a general network for 1 or multiple loops van also be extended to be applied to the other models that we studied in Sections SI 3 and SI 4. We refrain from giving those proofs here*.

## A Theorem of Petrovitch

The Theorem proved by Petrovitch [3] states that if *u* satisfies the differential inequality

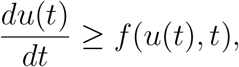

and *y* is the solution to the ODE

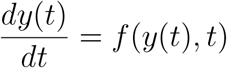

under the boundary condition *u*(*t*_0_) = *y*(*t*_0_), then: for *t* < *t*_0_, *u*(*t*) ≤ *y*(*t*) and *t* > *t*_0_, *u*(*t*) ≥ *y*(*t*)

